# Loss of Myosin-1e biases MMTV-PyMT induced breast cancer towards a differentiated and secretory state

**DOI:** 10.1101/2022.04.27.489205

**Authors:** Eric L. Plante, Sharon E. Chase, Ebbing P. de Jong, Theresa M. Curtis, Michael E. Garone, Mira Krendel

**Affiliations:** Department of Cell and Developmental Biology, SUNY Upstate Medical University, Syracuse, NY 13210, USA; Department of Biochemistry and Molecular Biology, SUNY Upstate Medical University, Syracuse, NY 13210, USA; Department of Biological Sciences, SUNY Cortland, Cortland, NY 13045, USA

## Abstract

Expression of the unconventional myosin, Myosin-1e (Myo1e), has been shown to contribute to tumor progression in the MMTV-PyMT mouse model of mammary tumorigenesis and is associated with poor outcome in breast cancer patients. However, the specific effects of Myo1e expression on the mammary tumor cells remain unidentified. Here, we used Myo1e-KO and wild-type (WT) MMTV-PyMT mice on a pure genetic background to further investigate the molecular and cellular effects of Myo1e expression. Myo1e-WT tumors were characterized by an increased abundance of intra-epithelial macrophages and lower amounts of the extracellular matrix. Transcriptomic profiling of WT and Myo1e-KO tumors identified a pattern of differential expression of tumor suppressor and tumor-promoting genes that was consistent with the observed differences in tumor progression and morphology between the genotypes, and also revealed differential expression of genes associated with secretion and cell-cell adhesion. In agreement with the RNA-seq findings, Myo1e-expressing tumor cells exhibited increased proliferation and elevated nuclear enrichment of YAP1 transcriptional activator compared to Myo1e-KO tumor cells. To investigate tumor cell-autonomous effects of Myo1e expression, we used the epithelial cell line PY-230 derived from the MMTV-PyMT-induced mouse tumor to create Myo1e-depleted cells by Crispr-mediated genome editing. Cells deficient in Myo1e had increased expression of genes encoding milk components compared to the wild-type cells. Electric cell-substrate impedance sensing (ECIS) measurements showed that depletion of Myo1e in PY-230 cells resulted in increased resistance to electrical current indicating enhanced epithelial barrier function. Overall, we find that Myo1e expression biases tumors towards a less-differentiated, pro-tumorigenic state, and that depletion of Myo1e is associated with a pro-secretory, more differentiated state.

## Introduction

Myosins are actin-dependent molecular motors that move along actin filaments and can modulate actin cytoskeletal organization. Some myosins have been linked to either promoting or suppressing malignant transformation (1). An unconventional myosin, Myosin 1e (Myo1e), has been identified as part of a gene signature associated with poor outcome in patients with basal-like breast cancer (2), and the expression of Myo1e was also found to be associated with poor patient survival in grade-1 breast cancer (3).Previously, our lab has studied the effects of Myo1e on breast cancer progression using the MMTV-PyMT model of tumorigenesis (3). In this model, the oncogene PyMT is expressed in mouse luminal epithelial cells to reproducibly induce breast tumors with subsequent metastasis to the lung (4). Knockout of Myo1e led to a more-benign tumor morphology with well-organized layers of epithelial cells lining enlarged ducts, decreased proliferation of tumor cells, and reduction in tumor metastasis (3). While this model allowed us to characterize tumor morphology, insight into the molecular effects of Myo1e on the tumors was limited, with one confounding factor being mixed genetic background (FVB and CL57BL/6) of the mice used for the initial study. Strain background (FVB vs. CL57BL/6) has significant effects on both tumor latency and growth rate in the MMTV-PyMT model (5). In order to elucidate the molecular and cellular differences between Myo1e WT and KO tumors, we backcrossed our Myo1e-KO and WT MMTV-PyMT mice to the CL57BL/6 mouse strain to obtain mice on a pure genetic background.

The MMTV-PyMT model is used extensively to study the effects of specific genes as well as the role of the tumor microenvironment and immune system on breast cancer tumor initiation and progression (4). Tumors arising from the PyMT oncogene expression resemble the luminal B subtype of human cancer (6), and MMTV-PyMT-driven tumor progression is reminiscent of the histological features of human breast cancer progression (7). Furthermore, changes in protein biomarkers and genetic alterations reveal further similarity between MMTV-PyMT mouse tumors and human cancer (4, 8). Overall, the MMTV-PyMT mouse is a useful model for the elucidation of the Myo1e-dependent features of tumor progression with relevance to human disease.

Recent studies have shown that Myo1e expression and activity is increased downstream of some pro-tumorigenic and cell motility associated signaling pathways. Specifically, Myo1e expression was increased following TGF-beta signaling (along with the unconventional myosin Myo10) in a model of lung cancer, and silencing of Myo1e reduced cancer cell motility in response to TGF-beta (9). Furthermore, Myo1e activity was increased downstream of ERK signaling to increase cancer cell motility (10). Myo1e expression was also found to increase in response to the activation of the Rankl signaling pathway in osteoclasts (11), and Rankl signaling is also known to be involved in breast cancer progression. Myo1e expression alters the activity of signaling proteins such as FAK, AKT, and Rac1 in lymphocytes (12), and FAK in melanoma cells (13). While some of the roles of Myo1e that may impact cancer progression have been studied, specific signaling pathways and cell types that are affected by Myo1e to change breast cancer progression and tumor morphology in the MMTV-PyMT model have yet to be identified. We therefore aimed to identify molecular and cellular features of the MMTV-PyMT driven tumor progression that are affected by the Myo1e knockout.

To identify possible tumor-promoting or tumor-suppressing genes affected by Myo1e, we performed transcriptomic analysis of Myo1e-WT and KO MMTV-PyMT tumors using RNA-seq and characterization of Myo1e-WT and KO tumors using immunohistochemistry. Tumors expressing Myo1e exhibited increased markers of proliferation, nuclear enrichment of the transcription co-factor YAP, and infiltration of intra-epithelial macrophages while tumors depleted of Myo1e were characterized by elevated expression of milk components and accumulation of milk-associated proteins in the tumor cystic fluid. Elevated expression of casein upon Myo1e depletion was also observed in PyMT-derived Py230 mammary tumor cells, indicating that Myo1e may regulate expression of milk-associated genes in a cell-autonomous manner. Interestingly, Myo1e expression also affected the paracellular junctional strength of the Py230 cells. Overall, our findings suggest that Myo1e expression promotes activation of pro-tumorigenic pathways in breast cancer cells while the depletion of Myo1e biases tumor cells towards a lactogenic differentiated state.

## Materials and Methods

### Animal Studies

All animal experiments were approved by the SUNY Upstate Medical University IACUC under animal protocol 373. Myosin-1e knockout mice have been previously described (14). Genotyping for Myo1e knockout allele and the MMTV-PyMT transgene was performed as described (3). MMTV-PyMT carrying Myo1e WT or Myo1e KO mice on the mixed FVB/CL57BL/6 background (3) were backcrossed to the CL57BL/6 strain mice for at least six generations. All animals used for experiments contained a single copy of the MMTV-PyMT transgene and were on the pure CL57BL/6 background. Tumor latency was determined using regular palpation, and growth rate measurements were performed using digital calipers as previously described (3).

### Cell culture

Py230 cells were purchased from the ATCC®(CRL-3279, RRID CVCL_AQ08). Py230 cells were grown in F-12 media with 5% FBS and Mito+ serum extender (Corning®). Crispr-Cas9 mediated depletion of Myo1e expression was performed by the Synthego Corporation (Cambridge, MA, USA). Briefly, a guide sgRNA:AAGGCGCCUACCGGUACCAU was used to target Crispr-Cas9 to exon 2 of the MYO1E gene in order to create premature stop-codons following insertions or deletions within the nucleotide sequence. Myo1e protein expression was measured via Western Blot analysis using rabbit polyclonal antibodies described previously (15), RRID AB_2909514. The Myo1e-KO cells were then utilized for experiments.

Primary tumor cells were extracted from tumors and grown on coverslips for immunostaining as described in (3). Primary tumor cells were cultured in 1:1 DMEM:F12 media with 2mM glutamine, 2% FBS and antibiotic/antimycotic solution.

### Antibodies

α-tubulin (mouse monoclonal, clone DM1α (Thermo Scientific, Waltham, MA)), E-Cadherin (mouse monoclonal (BD Transduction Laboratories, San Jose, CA)), Ki-67 (rabbit monoclonal, clone SP6 (Thermo Scientific, Waltham, MA)), Cyclin-D1 (rabbit polyclonal against C-terminus (MBL International, Woburn, MA)), Fibronectin (Polyclonal rabbit anti-Fibronectin, Sigma-Aldrich), YAP (rabbit monoclonal, Cell Signaling Technology, Danvers, MA), Anti-mouse-milk (Polyclonal rabbit, Nordic MUBio, Susteren, The Netherlands), Iba1 (Polyclonal rabbit, Wako Chemicals, Richmond, VA).

### Immunohistochemistry and Histology

Immunohistochemical staining (IHC) of cryosections: excised mammary tumors were fixed in 4% paraformaldehyde /phosphate buffered saline (PBS) solution for 1 hour prior to flash freezing in OCT and cryosectioning at 6 µm. Slides were rehydrated in PBS, blocked with a 3% BSA/PBS solution for 30 minutes, incubated with primary antibody against fibronectin in blocking buffer (3% BSA/PBS solution) for 2 hours and fluorescently labeled secondary antibody for 1 hour. Images were collected using Perkin Elmer Ultraview VoX Spinning Disc Confocal system mounted on a Nikon Eclipse Ti microscope, equipped with a NIKON Apo TIRF 60X 1.49NA oil objective and a Hamamatsu C9100-50 EMCCD camera and controlled by Volocity software.

IHC of formalin-fixed and paraffin-embedded (FFPE) tumor tissue: DAB based staining was performed using the Vectastain® Elite® ABC HRP kit (Vector Laboratories™, Inc, Burlingame, CA, USA). To do this, FFPE tissue was deparaffinized with xylene and rehydrated in a series of alcohol baths of absolute, 95% and 70% alcohol before being hydrated in water. Antigen retrieval was performed in a pressure cooker with 10mM NaCitrate (pH 6) for 7 minutes. The slides were placed in Tris-buffered saline (TBS) for 5 minutes, followed by a 45 minute blocking step using a 2.5% normal horse serum/TBS solution. When antibodies raised in mice were used, slides were blocked by mouse on mouse Ig blocking reagent from the M.O.M. ® Immunodetection Kit (Vector Laboratories™, Inc, Burlingame, CA, USA) using concentrations and incubation times specified in the kit. Primary antibody was also diluted in the M. O. M. protein diluent, and the M. O. M biotinylated antibody was used. The slides were then incubated at 37°C with primary antibodies diluted in blocking buffer. Primary antibodies used to stain FFPE tissue include antibodies to Myosin 1e, Fibronectin, Ki67, Cyclin-D1, Iba1, and Milk. The slides were then washed with TBS before being incubated at 37°C with secondary biotinylated antibody diluted in blocking buffer provided with the kit. The slides were washed again with TBS and then incubated with Vectastain® ABC reagent for 30 minutes. The slides were washed with TBS and incubated with ImmPACT HRP substrate reagent from the Vector® ImmPACT DAB substrate kit (Vector Laboratories™, Inc, Burlingame, CA, USA). The slides were then washed with water before being counterstained with hematoxylin and coverslipped.

H&E and Masson’s Trichrome stain was performed on formalin-fixed and paraffin embedded tissue. FFPE tissue was first de-paraffinized in xylene and hydrated in 95%, 70% alcohol then briefly dipped in water. Tissue was then stained with Hematoxylin for 5 minutes, dipped in water, decolorized in 0.5% acid alcohol, “blued” in ammonia water and washed under tap water. The tissue was then counterstained with Eosin for 15 seconds, dehydrated in alcohol, cleared in xylene and coverslipped. A similar process was followed for Masson’s Trichrome staining but Weigert’s hematoxylin was used and stained the tissue for 10 minutes. Following a water wash, the tissue was stained with Biebrich Scarlet for 2 minutes. The tissue was then put into a phosphotungstic/phosphomolybdic acid solution for 10 minutes followed by aniline blue for 5 minutes, a water rinse and wash with 1% acetic acid solution for 5 minutes. Finally, the tissue was rinsed with water, dehydrated, cleared in xylene and coverslipped. Histological and immunostained slides were imaged using an Olympus CHBS microscope and Canon Rebel T3i EOS 600D camera.

### Image analysis of IHC samples

Analysis of proliferation index using Ki-67 staining was performed using 3 age-matched mice of each genotype. Briefly, tumors from 18-week-old mice were sectioned stained for Ki-67. The number of Ki-67 positive tumor cell nuclei and total number of tumor cell nuclei (labeled with hematoxylin) was manually counted from 5 fields of view per animal. Analysis of Cyclin-D1 staining was performed on 3 age-matched mice of each genotype. Tumor sections were stained for Cyclin-D1 from 18-week-old mice. The number of Cyclin D1-positive peripheral tumor cells and non-peripheral Cyclin D1-positive tumor cells was determined for 5 tumor acini per animal from 5 fields of view. A tumor cell was considered peripheral if it resided on the boundary between the tumor acinus and the stroma. The fraction of peripheral Cyclin-D1 positive tumor cells to total tumor cells was then calculated.

Analysis of Iba1 staining was performed on 3 age-matched mice of each genotype. Tumor sections from 18-week-old mice were stained for Iba1. The number of intra-epithelial Iba1 positive cells within tumor acini was then counted from 5 fields of view per animal.

### Quantification of fibronectin area

At least 6 fields of view per tumor from 3 Myo1e WT and 3 Myo1e KO MMTV-PyMT mice were analyzed. Brightness and contrast were standardized, and pixels containing stroma-associated fibronectin signal were manually segmented in ImageJ to measure fibronectin-positive area. To measure the overall tissue area (excluding the acellular background), pixel intensity threshold was used to create a mask (Fig.1i). The percent area of fibronectin was calculated as the number of pixels labeled as 1 (true) in the binary manually segmented mask of fibronectin divided by the number of pixels corresponding to the tissue mask.

### Immunofluorescence staining of primary tumor cells

Analysis of Cyclin-D1 expression in primary tumor cells was performed using 3 Myo1e-WT and 3 Myo1e-KO MMTV-PyMT mice. Primary tumor cells were extracted and stained for Cyclin-D1. Briefly, tumor cells plated on coverslips and were cultured for 48 hours before being fixed in 4% PFA/PBS for 15 min, washed with PBS, permeabilized using 0.2% Triton-X/PBS and blocked using 3% BSA/PBS. The tumor cells were then stained with primary antibody diluted in blocking buffer for 1 hour, washed, and stained with secondary antibody for 1 hour. Images were captured using a Perkin Elmer Ultraview VoX Spinning Disc Confocal system mounted on a Nikon Eclipse Ti microscope and equipped with a NIKON Apo TIRF 60X 1.49NA oil objective and a Hamamatsu C9100-50 EMCCD camera controlled by Volocity software. 7 fields of view were analyzed per animal. The total number of cells per field of view was assessed using DAPI stain, which was applied in the solution with the secondary antibody. The number of Cyclin-D1 positive nuclei was then counted per field of view. A ratio of Cyclin-D1 positive nuclei to total nuclei was then calculated per field of view. Analysis of YAP staining of primary tumor cells was performed using cells from 4 Myo1e-WT and 4 Myo1e-KO MMTV-PyMT mice. Staining of primary tumor cells for YAP was performed as described above for Cyclin-D1. To quantify nuclear enrichment, rectangular ROIs (44x40 pixels) were used to measure mean fluorescence intensity within a region of nuclei (determined by DAPI staining) and adjacent cytosol. The ratio of nuclear to cytoplasmic mean fluorescence intensity was then calculated for at least 90 cells per genotype.

### Analysis of cystic fluid and protein identification

To test for the presence of proteinaceous secretions in Myo1e-KO MMTV-PyMT tumors, cystic fluid from eight KO tumors was collected by aspiration from tumors during dissection using a P1000 micropipette. Samples were boiled with SDS-PAGE sample buffer, separated on an SDS-page gel, and stained with Coomassie Blue. The amount of protein in the cystic fluid (whole-lane intensity) was calculated by running samples with BSA standards of known concentrations to generate a standard curve. 100 µg of protein (a representative sample from a single tumor) was submitted for MS/MS protein identification. Using the FASP method (16), a mixture of Tris-HCl pH 8.5, SDS and DTT were added to the sample to a final concentration of 10 mM, 0.4% (w/w) and 10 mM. Disulfides were reduced by heating to 95°C for 5 min, and the samples were cooled to room temperature before being transferred to a 10 kDa MWCO ultrafiltration vessel (Pall, OD010C33). 200 µl of 8M urea containing 100 mM Tris pH 8.5 was added, and the filters were centrifuged to near-dryness. Cysteine residues were alkylated with 50 mM iodoacetamide (IAM) for 25 min in the dark, followed by centrifugation. The retained proteins were further washed with three aliquots of urea + Tris, followed by three washes of 50 mM ammonium bicarbonate. Trypsin, in 40 µL of 50 mM ammonium bicarbonate, was added at a ratio of 0.02 µg per µg of protein sample and digestion was allowed to proceed overnight at 37°C. The resulting peptides were collected in a clean tube by centrifugation, followed by a wash with 50 µL of 0.5 M NaCl. The peptides were desalted using mixed-mode cation exchange (MCX) stage tips (17). The 200 µL tips were packed with two cores of Empore MCX material (SDB-RPS, 3M, 2241) made using 14-gauge blunt needles. The sorbent was conditioned with acetonitrile (ACN), washed with solvent A (3% ACN in water with 0.2% TFA), up to 10 µg of sample was loaded, washed twice with solvent A, washed once with solvent B (65% ACN in water with 0.1% TFA) and eluted using 65% ACN in water containing 5% (v/v) of ammonium hydroxide. The desalted peptides were dried in a speed-vac. LC-MS analysis was performed using a Waters nanoAcquity LC and autosampler coupled to an Orbitrap XL hybrid ion trap Orbitrap mass spectrometer. Two µL of sample was injected onto a reversed phase nano LC column from New Objective (PicoFrit, 75 µm I.D. with 15 µm tip packed with 10 cm of BioBasic C18 particles, 5 µm particles). A linear LC gradient from 2 to 35% ACN over 45 min with constant 0.1% formic acid was used, followed by a short ramp to 90% ACN and re-equilibration to 2% ACN for a total run time of 60 min. The LTQ-Orbitrap was operated in a top-five data-dependent mode using survey scans at 30,000 resolution from 375-1800 m/z. Tandem MS scans were acquired in the ion trap with an isolation width of 2 m/z and fragmentation mode was CID with 35% normalized collision energy for 0.1 ms. The automatic gain control settings were 3×10^5^ ions in the ion trap, and 1×10^6^ in the Orbitrap. Dynamic exclusion was used with a duration of 15 s and a repeat count of 1. Tandem mass spectra were searched against a mouse database (Uniprot, 51444 entries), and common contaminant proteins using the Sequest HT node in Proteome Discoverer 1.4. Search parameters included variable modification of methionine oxidation, and partial trypsin specificity with 2 missed cleavages. Search results were filtered to Percolator q-values < 0.01 and peptide confidence filter set for “high”.

### RNA sequencing

Tumor samples used for RNA-seq were immediately flash frozen in liquid nitrogen following excision and stored at -80°C. RNA was extracted using a TRIzol™-based protocol. 500ng of RNA from each sample was used for Illumina Truseq Stranded mRNA library Prep Kit. An Agilent 2100 bioanalyzer with DNA 1000 chip was used to validate library size. The libraries were then quantified on a Qubit 3.0 fluorometer. Single end, 1x75bp sequencing was performed on an Illumina NextSeq^®^ 500 system (Illumina, inc. San Diego, CA) utilizing a High-Output kit. Sequence processing, including read trimming, BowTie2 alignment to the mm10 reference genome and quantification via counts per million was done using Partek® Flow software. P-value was determined using an Akaike-defined multimodel approach (normal, lognormal, negative binomial or Poisson). This profiling yielded 79 differentially expressed genes beyond +/-1.5 fold expression change and a relative expression p-value of p<0.01 (S1 Table). The data was visualized using principal component and t-SNE analysis to reduce the dimensionality of the data. Coloring of the data points by genotype reveals that Myo1e KO and WT cluster distinctly in this reduced space (S1 Fig).

### qRT-PCR of milk-associated genes

To generate a lactogenic response in Py230 cells, 10 U/mL Prolactin (Sigma-Aldrich®, St. Louis, MO, USA) and 100 µM Dexamethasone (Sigma-Aldrich®, St. Louis, MO, USA) were added to growth media for 48 hours. Total RNA was extracted using the RNeasy Mini kit from Qiagen® (Qiagen inc.), and cDNA was synthesized using the iScript cDNA synthesis kit by Qiagen® (Qiagen inc.). 200ng of cDNA from each sample was used to perform qRT-PCR using Bio-Rad® iTaq universal SYBR Green Supermix (Bio-Rad Laboratories, Inc., USA) on a Bio-Rad® CFX 384 Real Time PCR System (Bio-Rad Laboratories, Inc., USA). Primers: fGapdh:ACTCCACTCACGGCAAATTC, rGapdh:TCTCCATGGTGGTGAAGACA, fRplp0:GTTCTCCTATAAAAGGCACAC, rRplp0:AAAGTTGGATGATCTTGAGG, fPigr:AAGAACTCCAGAGATTTGGG, rPigr:GTGGTAGTCACGATTTCATC, fCsn2:GTATTTCCAGTGAGGAATCTG, rCsn2:ATTGCAAGAGATGGTTTGAG, fCsn3:CTTTTTGGCTGCAGAGATAC, rCSn3:AAGTGGGTAGTACATGTATGG. Calculation of relative expression was performed using the ΔΔCt method.

### Transfection of Py230 cells

Myo1e-KO Py230 cells were utilized for transfection with C1-mEmerald-Myo1e using Lipofectamine 3000 ^TM^ reagent (Thermo Scientific, Waltham, MA). Cells were cultured on coverslips in a 12-well plate. The Lipofectamine 3000 ^TM^ reagent manufacturer protocol was used with 1 ug of plasmid DNA per well. 24 hours post-transfection, the transfected cells were fixed, stained using anti-E-cadherin and imaged as described above for immunofluorescence assays.

### Electric cell-substrate impedance sensing

To acquire complex impedance data, Electrical cell-substrate impedance sensing was used (ECIS^TM^, Applied Biophysics Inc., Troy, NY, USA). A Z-Theta instrument was used with a 96 W array station. 20,000 Py230 cells per well were seeded into a 96W10idf PET array pre-coated with collagen I and grown to confluence over 3 days. ECIS complex impedance data, which were comprised of cell barrier resistance (R) and membrane capacitance (C), were obtained using a multiple frequency time (MFT) course setting ranging from 62.5 to 64,000 Hz. Through ECIS software modeling, the R was further broken down into Rb (resistance caused by paracellular junctions) and α (resistance caused by cell-substrate junctions).

## Results

### Myo1e influences tumor organization and ECM accumulation in the tumor stroma

We analyzed mammary tumor growth in the MMTV-PyMT mice on a pure C57BL/6J genetic background either expressing (WT) or depleted (KO) of Myo1e. Western blot analysis confirmed the expression of Myo1e in the Myo1e-WT CL57BL/6 mouse tumors and lack of Myo1e in the Myo1e-KO CL57BL/6 mouse tumors (Fig 1a). Concordant with the previous observations made using mice with a mixed C57BL/6J and FVB genetic background (3), on gross examination the tumors lacking Myo1e were cystic and filled with fluid, while tumors from the Myo1e-expressing mice were more solid (Fig 1b). Histological examination of tumors from age-matched mice (18 weeks old) showed that tumors from WT mice consisted of solid masses of poorly differentiated tumor cells, while tumors from the Myo1e-KO mice maintained duct-like tissue architecture (Fig 1c). Specifically, Myo1e KO tumors contained expanded mammary ducts, abundant acellular/fluid-filled areas, and fibrovascular stalks protruding into the lumens of the enlarged ductal structures. This difference in tumor morphology was similar to that observed in our earlier study (3).

**Figure 1.**
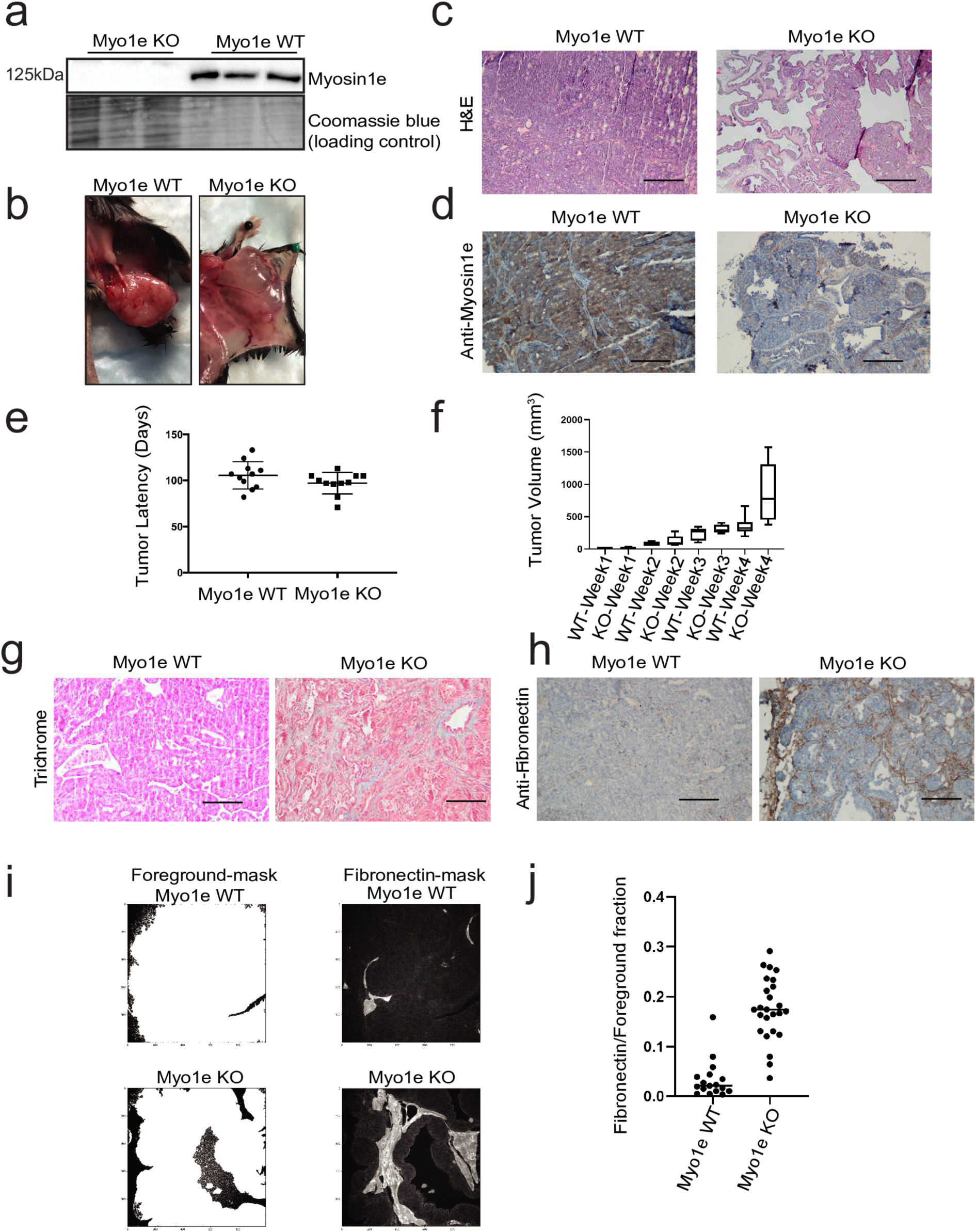
Myo1e coordinates a malignant tumor growth pattern. (a) Western Blot showing loss of Myo1e expression in tumor lysates of Myo1e KO mice. Gross examination of tumors (b) and H&E staining of FFPE age-matched tumor tissue (c) shows that Myo1e-KO masses are cystic and fluid-filled. (d) Immunostaining of age-matched MMTV-PyMT tumors using anti-Myo1e, counterstained with hematoxylin, showing Myo1e expression in tumor cells. (e) Tumor latency in Myo1e-WT and -KO mice. Each data point represents mouse age (in days) until the appearance of a palpable tumor. (f) Growth of Myo1e-WT and -KO tumors (expressed as the volume of the tumor). (g) Masson’s Trichrome staining of FFPE tumor tissue. (h) Immunostaining of age-matched MMTV-PyMT tumors using antibody to fibronectin, counterstained with hematoxylin. (i) Analysis of the fraction of tumor tissue that contains stromal fibronectin. To create a foreground mask, a pixel intensity threshold was used to identify foreground (tissue) from background (acellular area) in the images of tumor tissue. Fibronectin was manually segmented using ImageJ software (Fibronectin mask) and the area of tissue that contains fibronectin was then calculated. (j) Quantification of the fraction of tumor tissue that contains fibronectin, each data point represents a field of view. Statistical analysis was performed using Student’s t-test. ***p<0.0001. Scale bars: (c) 250µm, (d, g, h) 100µm.

Immunohistochemical staining with anti-Myo1e antibody showed that Myo1e was enriched in tumor cells compared to stromal cells (Fig 1d). Unlike our previous observations, which showed on average a 19-day increase in tumor latency in the Myo1e-KO mice on the mixed genetic background (3), no difference in tumor latency was detected in the current study (Fig 1e). Mice on the mixed genetic background had an average tumor latency of 75 days for Myo1e-KO tumors and 56 days for Myo1e-WT tumors (3). In comparison, Myo1e-KO tumors on a C57BL/6J genetic background have an average tumor latency of 97 days and Myo1e-WT tumors have an average latency of 101 days, with no statistically significant differences between the two groups (Fig 1e). Volume of the Myo1e-KO tumors increased faster than that of Myo1e WT tumors, likely due to fluid accumulation (Fig 1f). Longer tumor latency on the C57BL/6 is consistent with the previous findings from other groups (5).

Proteins involved in cell-ECM adhesion, such as fibronectin and integrins, are associated with mammary gland function, mammary epithelial cell shape, expression of secretory genes, and tumor behavior (18–24). We investigated if differences in the ECM composition are present between the Myo1e-WT and KO tumors. Masson’s Trichrome staining revealed that the stroma/fibrovascular stalks in KO tumors were enriched in ECM (Fig 1g). Immunostaining against fibronectin showed that Myo1e-KO tumors contained more stromal fibronectin compared to age-matched Myo1e-WT tumors (Fig 1h). Quantification of anti-fibronectin staining revealed that fibronectin comprised 17.2% of KO tumor tissue (primarily in the fibrovascular stalks) compared to the 5.8% of WT tumor tissue, confirming that Myo1e-KO tumors are enriched in ECM compared to Myo1e-WT tumors (Fig 1i and 1j). Overall, we find that Myo1e expression in MMTV-PyMT mice on a CL57BL/6J genetic background contributes to rapid tumor progression towards the loss of mammary tissue architecture, while tumors that do not express Myo1e progress slower and maintain ductal organization with enlarged, fluid-filled lumens.

### Differential expression of tumor-promoting and -suppressing genes in the Myo1e-WT and KO tumors

Transcriptomic examination of differentially expressed genes has previously uncovered biomarkers that are involved in tumor progression and metastasis in the MMTV-PyMT model (25). Given that tumor latency in the Myo1e-WT and KO mice on the pure genetic background was similar, we used tumors from age-matched Myo1e-WT and KO mice to compare gene expression profiles using RNA sequencing. Five WT and five KO tumor tissue samples from 18-week-old mice were used for RNA sequencing. Single end, 1x75bp sequencing resulted in an average of 24 million reads per sample. Following read processing, further described in methods, quantification using counts per million was determined. Differences in gene expression between genotypes were considered significant if expression change was more than 1.5-fold and relative expression p-value was less than 0.01. Seventy-nine genes were identified as significantly differentially expressed between the WT and KO tumor samples (Fig 2 and S2 Fig). The top 20 differentially expressed genes (DEGs) between the genotypes are displayed in Fig 2b. Some DEGs identified by the RNA-seq analysis have been previously implicated in tumor growth, metastasis, and invasion, as further described in the Discussion.

**Figure 2.**
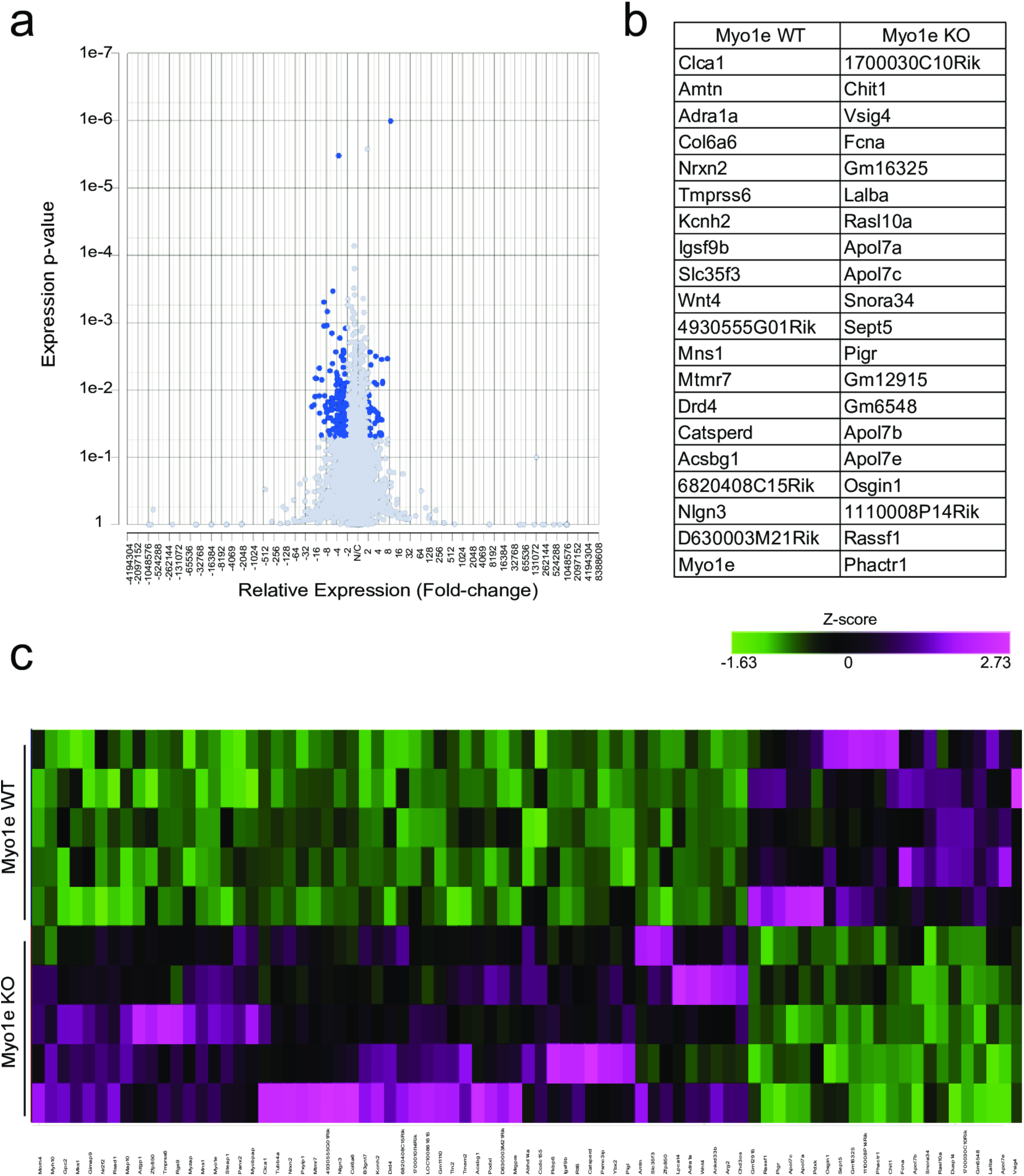
Differential gene expression in the Myo1e-WT and KO tumors. (a) Volcano plot displaying the distribution of differentially expressed genes (DEGs) between WT and KO tumors. Highlighted DEGs meet the criteria of being differentially expressed at least 2-fold and have a p-value range of 0.000001-0.05. (b) The most differentially expressed genes expressed in five Myo1e KO tumors (right column) and five Myo1e WT age-matched tumors (left column). The displayed genes are those with the top 20 largest expression fold-change in the assigned genotype that have an expression p-value of <0.01. (c) Heat map displaying Z-scores for 79 differentially expressed genes (beyond +/-1.5-fold expression change and a relative expression p-value of p<0.01).

Transcripts associated with increased proliferation and cell migration were enriched in the Myo1e-WT tumors while those negatively associated with cancer cell proliferation were enriched in the Myo1e-KO tumors. In addition to the tumor-suppressing genes, we found significant upregulation of genes associated with mammary cell secretory differentiation in Myo1e-KO tumors, including Lalba, Pigr, Osgin1 and Phactr1. Expression of these genes is increased during lactation (26–30). Overall, the transcriptomic analysis of Myo1e WT and KO tumors identified pro-tumorigenic genes expressed in Myo1e-WT tumors, and tumor-suppressive and differentiation-associated genes expressed in Myo1e-KO tumors.

Using the NetworkAnalyst 3.0 analytical platform (31), we performed Gene Ontology (GO) gene enrichment analysis of the seventy-nine differentially expressed genes based on the PANTHER classification system. A cutoff p-value of p<0.05 was used to identify cellular compartments (Table 1) or biological processes (Table 2) that were characterized by significant enrichment of genes in our dataset. The highest degree of enrichment was found for the genes classified as associated with the extracellular region of the cell (Table 1) and cell adhesion (Table 2).

**Table 1.**
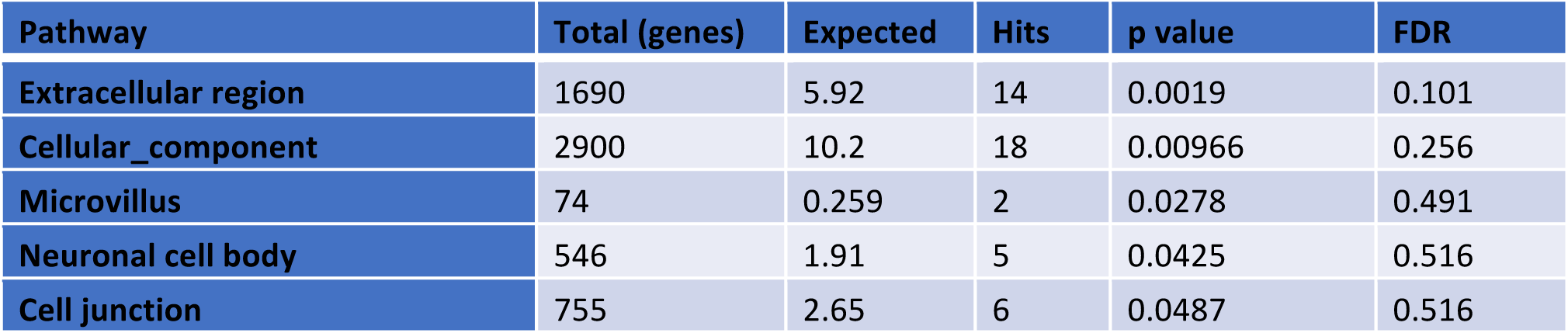
GO-Slim term enrichment analysis using the PANTHER: Cellular Compartments categories. Significantly differentially expressed genes in the RNA-seq data set were compared to reference lists of genes of GO-Slim terms of the PANTHER: Cellular Compartments categories via the NetworkAnalyst 3.0 analytical platform (31). The number of “Hits” are the genes within the RNA-seq data set that are statistically enriched when compared to the total gene reference list for each category. The number of statistically “Expected” genes within a gene set compared to the reference list is displayed. A hypergeometric test was used to compute p-values and the false discovery rate (FDR) correction is shown.

**Table 2.** GO-Slim term enrichment analysis using the PANTHER: Biological Processes categories. Significantly differentially expressed genes in the RNA-seq data set were compared to reference lists of genes of GO-Slim terms of the PANTHER: Biological Processes categories via the NetworkAnalyst 3.0 analytical platform (31). The number of “Hits” are the genes within the RNA-seq data set that are statistically enriched when compared to the total gene reference list for each category. The number of statistically “Expected” genes within a gene set compared to the reference list is displayed. A hypergeometric test was used to compute p-values and the false discovery rate (FDR) correction is shown.

| Pathway | Total (genes) | Expected | Hits | p value | FDR |
| --- | --- | --- | --- | --- | --- |
| Cell adhesion | 516 | 1.77 | 9 | 5.51E-05 | *0.0106 |
| Lipid transport | 124 | 0.425 | 4 | 0.000847 | 0.0817 |
| Cell-cell signaling | 61 | 0.209 | 2 | 0.0185 | 0.927 |
| Chemical synaptic transmission | 166 | 0.569 | 3 | 0.0192 | 0.927 |
| Spermatogenesis | 489 | 1.68 | 5 | 0.0255 | 0.983 |

### Myo1e expression is associated with tumor cell proliferation and nuclear enrichment of the transcriptional coactivator YAP in primary tumor cells

Differential expression of tumor-suppressive and pro-tumorigenic genes between Myo1e-WT and KO tumors led us to test whether there were differences in tumor cell proliferation between the genotypes. Sections of age-matched Myo1e-WT and KO tumors were immunostained for the proliferation markers Ki67 and Cyclin D1 (Fig 3a and 3b). Myo1e-WT tumors contained a larger fraction of tumor cells that were Ki67-positive compared to Myo1e-KO tumors (Fig 3c). Progression of PyMT-induced tumorigenesis is associated with a transition from the Cyclin D1 expression primarily in peripheral cells within mammary tumor acini to a more uniform expression of Cyclin D1 throughout the tumors [7]. Cyclin D1 staining revealed that a larger fraction of Cyclin D1-positive cells resided at the periphery of tumor acini of Myo1e-KO tumors when compared to age-matched Myo1e WT tumors, indicating a less-advanced stage of tumor progression in the Myo1e-KO mice (Fig 3d). To test if Myo1e’s effect on tumor cell proliferation was also evident in tumor cells outside of the tumor milieu, primary tumor cells were extracted from WT and KO mice and examined *ex-vivo* (Fig 3e). Analysis of Cyclin D1 staining of primary tumor cells revealed that Myo1e WT cells contained a higher fraction of Cyclin D1-positive nuclei when compared to Myo1e KO primary tumor cells (Fig 3f).

**Figure 3.**
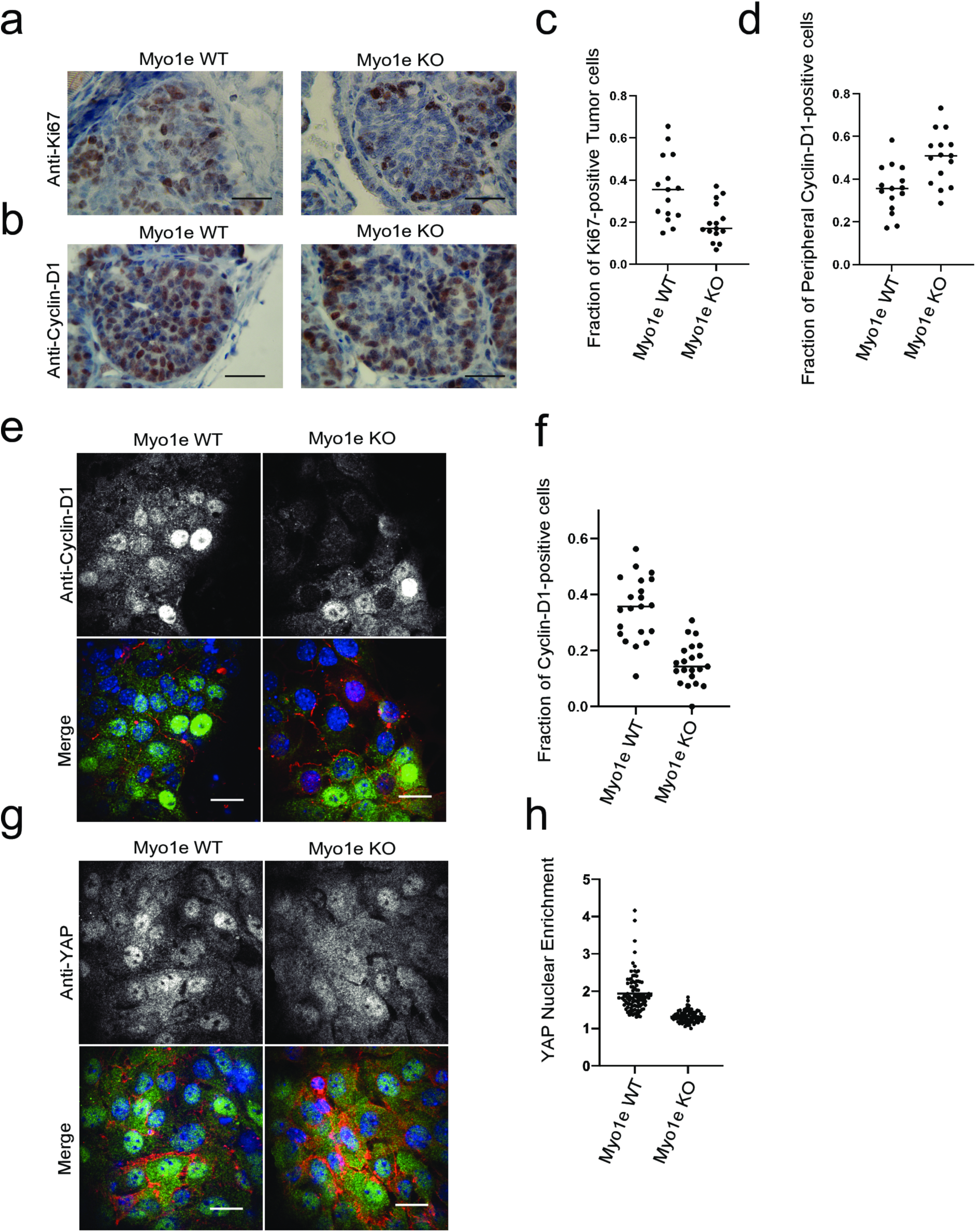
Myo1e expression is associated with tumor cell proliferation and nuclear enrichment of the transcriptional coactivator YAP in primary tumor cells. (a, b) Immunostaining (brown) for Ki-67 (a) or Cyclin D1 (b) with hematoxylin counterstain in age-matched 18-week-old Myo1e WT and KO mice. (c) Quantification of the fraction of Ki-67 positive tumor cells (out of all cells) per field of view in Myo1e-WT and KO tumors, p<0.005. (d) Quantification of the fraction of peripheral Cyclin D1-positive tumor cells (out of all Cyclin D1-positive cells) in Myo1e-WT and KO tumors (see Methods for the description of the quantification), p<0.005. Three mice per genotype and 5 fields of view were used for the quantification (c, d). (e) Immunostaining for Cyclin-D1 (green) and E-cadherin (red) in primary tumor cells from Myo1e-WT and KO mice. (f) Quantification of the fraction of Cyclin D1-positive tumor cell nuclei per field of view, p<0.001. Nuclear DAPI staining is shown in blue. Three mice of each genotype were used for cell extraction, and at least 475 cells per genotype were analyzed. (g) Immunostaining for YAP1 (green) and E-cadherin (red) in primary tumor cells from Myo1e-WT and KO mice. Nuclear DAPI staining is shown in blue. (h) Quantification of YAP1 nuclear enrichment for primary Myo1e-WT and KO tumor cells, p<0.001 (see Methods for the description of the quantification). Student’s T-Test was utilized for statistical analysis. Four mice of each genotype were used, and at least 90 cells per genotype were analyzed. Scale bars: (a, b) 25µm, (e, g) 20µm.

Gene targets of the cell growth-associated Hippo pathway were identified as DEGs in RNA-seq analysis of Myo1e WT and KO tumors. Specifically, Ingenuity pathway analysis (IPA, Qiagen Inc., https://www.qiagenbioinformatics.com/products/ingenuity-pathway-analysis) was used to identify likely upstream master regulators of the gene expression profiles generated from our RNA-seq experiment using Causal Network Analysis (S2 Table). YAP1 was identified as one of twenty-one transcriptional regulators with a network bias-corrected p-value of p<0.05 (S2 Table). Six potential molecular targets of YAP1 were identified by the IPA analysis in the gene expression profiles of RNA-seq (ARG2,KCNH2,LALBA, MYH10,NR2F2,WNT4 (S2 Table)). Hippo signaling has been associated with hallmarks of cancer in addition to oncogenesis in the MMTV-PyMT mouse model (32–34). Moreover, a top DEG found in RNA-seq analysis, RASSF1, is associated with tumor suppression and coordination of the YAP/TAZ transcriptional coactivators (33, 35). Three of the top five master regulators identified by the IPA analysis potentially target RASSF1 (S2 Table). Given differences in cell proliferation and in expression of RASSF1 between the genotypes, we hypothesized that Myo1e expression may affect YAP activity. To test this, primary tumors cells were stained for the transcriptional coactivator YAP1 (Fig 3g). Quantification of YAP1 nuclear enrichment revealed that YAP1 was more enriched in the nuclei of Myo1e-expressing primary tumor cells compared to the Myo1e-KO primary tumor cells (Fig 3h). Overall, this suggests that expression of Myo1e is associated with both increased tumor cell growth and YAP1 nuclear enrichment.

### Myo1e-expressing tumors contain more intra-epithelial macrophages compared to tumors lacking Myo1e expression

Tumor-associated macrophages have been shown to promote tumor growth and invasion in the MMTV-PyMT model (36). RNA-sequencing analysis of the Myo1e-WT and KO tumors revealed differential expression of multiple transcripts that are known to be expressed in macrophages including Igsf9b (37), Dusp4 (38), Vsig4 (39), Arg2 (40). In addition, both Myo1e and its homolog Myo1f are highly expressed in macrophages and regulate their activity and functions (41, 42) . We hypothesized that the abundance of tumor-associated macrophages may be dependent on the presence of Myo1e. To test this, age-matched tumors were stained for the macrophage marker Iba1, and the number of Iba1-positive cells was assessed in tumors of both genotypes (Fig 4). Iba1-positive cells were present in the stroma of both tumor genotypes but appeared more abundant in the stroma of Myo1e-KO tumors compared to the Myo1e-WT tumors (Fig 4a). Conversely, the number of Iba1-positive cells that resided among the tumor epithelial cells was higher in Myo1e-WT tumors compared to Myo1e-KO tumors (Fig 4b and 4c). These results indicate that Myo1e expression affects the abundance of intra-epithelial macrophages.

**Figure 4.**
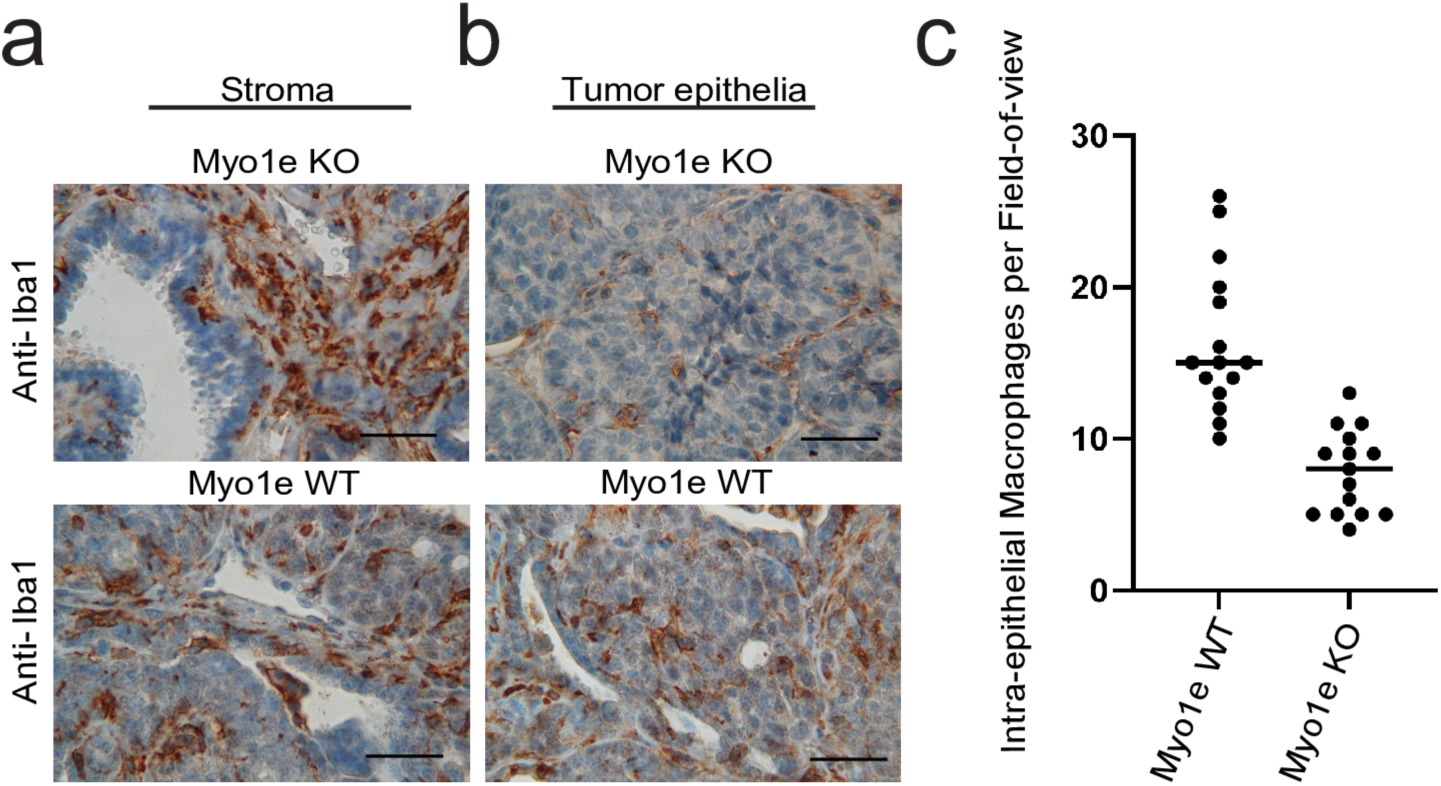
Myo1e-expressing tumors contain more intra-epithelial macrophages compared to the tumors lacking Myo1e expression. (a, b) Immunostaining for macrophage marker Iba1 (brown) with hematoxylin counterstain of the stroma (a) and tumor epithelium (b) of age-matched Myo1e-WT and KO tumor sections. (c) Quantification of the number of intra-epithelial Iba1 positive cells within tumor acini, p<0.001. Two-tailed student’s T-Test was utilized for statistical analysis. Five fields of view per animal from three age-matched mice of each genotype were used for quantification. Scale bars: (a, b) 25µm.

### Myo1e-KO tumors contain fluid enriched in milk-associated proteins

The depletion of Myo1e led to a transcriptional profile characterized by expression of secreted proteins (Table 1), and Myo1e-KO tumors were often fluid-filled (Fig 1). We therefore examined Myo1e-KO tumors for expression of milk-associated proteins. Differentiation of PyMT tumors towards expression of lactation-associated proteins has been previously described in this model. Specifically, in earlier studies in the PyMT model, blockade of Rank or Src signaling or depletion of the long non-coding RNA Malat1 led to the increased abundance of milk-associated proteins and expression of lactation-associated genes (43–45). SDS-PAGE analysis of cystic fluid aspirated from Myo1e-KO tumors showed a consistent band pattern across 8 cystic fluid samples from 8 KO mice, suggesting that similar proteins may be present in all samples (Fig 5a). We performed a proteomic analysis to characterize the proteins contained in the cystic fluid found in Myo1e-KO tumors. MS/MS analysis of the aspirate revealed that the majority of the top 10 proteins are constituents of milk, including casein (one of the major milk components) and serum proteins albumin and transferrin that are also found in milk (Fig 5b). We further examined the expression of milk proteins in cystic fluid using Western blot analysis. To do this, an antibody raised against pooled mouse milk was used for immunoblotting of aspirated cystic fluid protein lysates (S2 Fig). The Western blot analysis for mouse milk proteins generated a unique signal/band pattern in cystic fluid aspirate compared to WT tumor lysate samples (S2 Fig). We also examined milk protein expression in tumor tissue sections from Myo1e-WT and KO mice by immunohistochemical staining with anti-mouse milk antibodies. Anti-mouse-milk-protein staining was found in the apical regions of cells lining the enlarged ducts of Myo1e KO tumors (Fig 5c). Conversely, age-matched Myo1e-WT tumors were largely undifferentiated and showed little anti-milk staining (Fig 5c). Overall, it appears that depletion of Myo1e promotes expression of milk-associated proteins.

**Figure 5.**
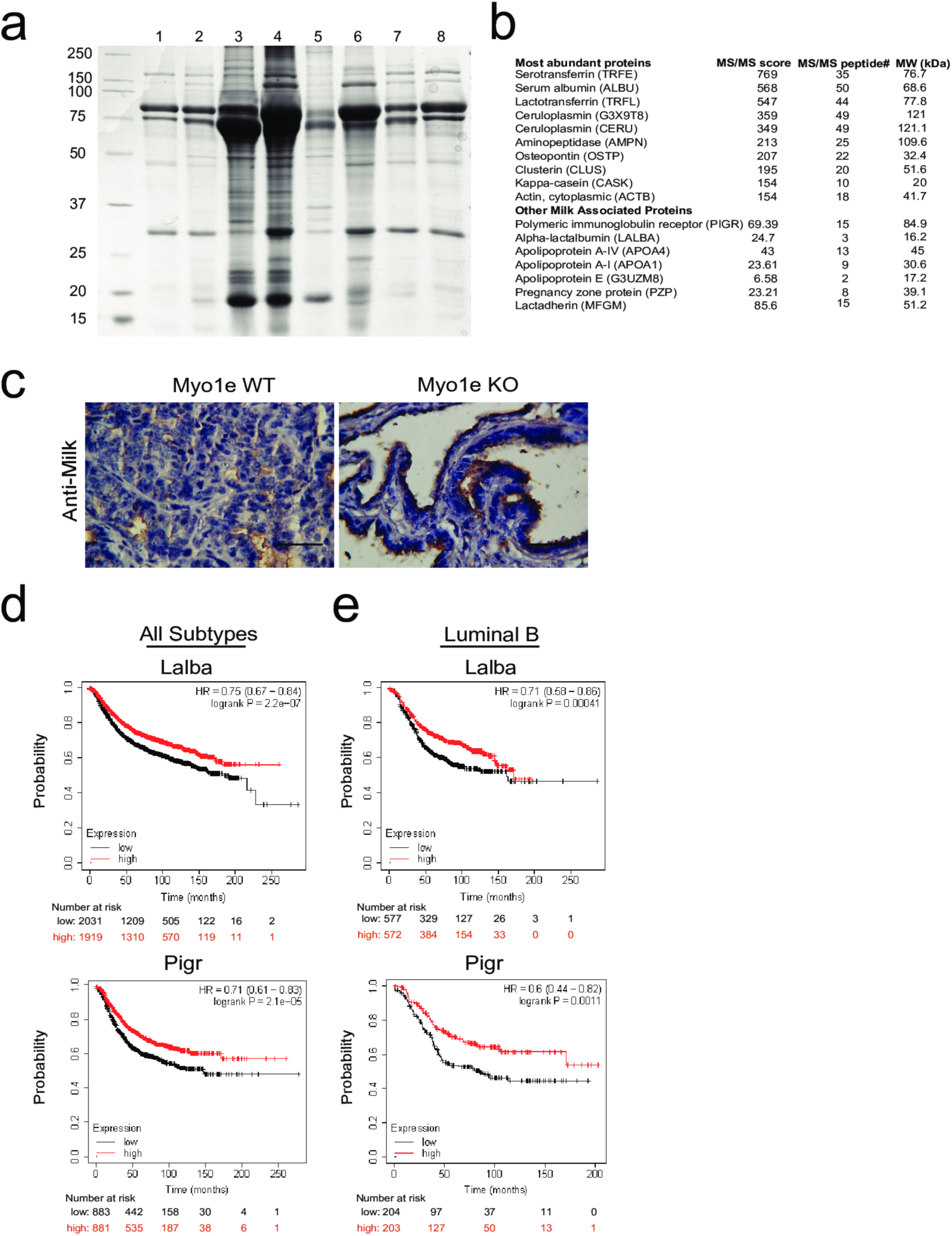
Myo1e-KO tumors contain fluid enriched in milk-associated proteins. (a) Coomassie-stained SDS-PAGE gel of aspirated cystic fluid from Myo1e KO tumors. (b) List of proteins identified in the Myo1e-KO cystic fluid by MS/MS. (c) Immunostaining of epithelial tumor tissue of age-matched Myo1e-WT and KO tumor sections with anti-mouse-milk antibodies (brown) with hematoxylin counterstain. (d) The probability of RFS over time in patients with all subtypes of breast cancer and expressing high or low abundance of Pigr or Lalba was analyzed using KM-plotter. (e) The probability of RFS over time in patients with the Luminal B subtype of breast cancer and expressing high or low abundance of Pigr or Lalba was analyzed using KM-plotter. HR= hazard ratio (with shown confidence interval). Statistical significance was calculated via Log rank P value. Scale bars: (c) 25µm.

Myo1e-KO luminal tumor cells lining the enlarged ductal structures appeared more columnar in shape compared to the more cuboidal-appearing luminal tumor cells in WT tumor acini. We measured the length of the nuclei along the basal to apical axis in luminal cells lining the ducts of Myo1e-WT and KO tumors and found that nuclei of KO cells were elongated compared to nuclei of WT luminal cells (S3 Fig). The Myo1e-KO luminal tumor cell nuclei elongation may be associated with their columnar-like cell shape.

We then tested if the expression of secretion-associated genes found in our RNA-seq analysis was relevant to human breast cancer patient outcome. We found that the expression of *Pigr* and *Lalba*, two genes generally associated with mammary cell secretory differentiation and upregulated in the more differentiated Myo1e KO tumors in our study, correlated significantly with human breast cancer patient outcomes identified using the KM-plotter (Fig 5d-5e)(46). Specifically, we found that the expression of *PIGR* and *LALBA* significantly correlated with relapse-free survival in all subtypes of breast cancer as well as specifically in luminal-B type breast cancer (which the MMTV-PyMT mouse model is thought to emulate).

### Py230 cells depleted of Myo1e have elevated milk-gene expression

To explore if Myo1e expression has a cell-autonomous effect on breast cancer cells (outside of the tumor/*in-vivo* context), we utilized a commercially available breast cancer cell line derived from the CL57BL/6 MMTV-PyMT mammary tumors, Py230 (47). These cells maintain the ability to form luminal-like tumors as well as display features of mammary cell differentiation upon lactogenic hormone treatment, including expression of the milk-associated transcript Beta-casein and formation of dome-like structures (47). We tested if Myo1e affects expression of lactation-associated genes in Py230 cells. Myo1e-KO Py230 cells were generated using Crispr-Cas9 (Fig 6a), and expression of lactation-associated transcripts was measured after treatment with lactogenic hormones prolactin and dexamethasone (47). Upon hormone treatment, both WT and Myo1e-KO Py230 cells appeared to expand in surface area compared to control cells (Fig 6b). Relative gene expression between Myo1e-WT and KO cells was measured using qRT-PCR of the lactation-associated genes Beta-casein (Csn2), Kappa-casein (Csn3), and the Polymeric immunoglobulin receptor (Pigr). Py230 cells depleted of Myo1e had significantly higher Csn2 and Csn3 expression, but not Pigr expression, when compared to Myo1e-WT Py230 cells following lactogenic hormone treatment (Fig 6c). Thus, Myo1e influences the expression of some lactation-associated genes in a cell-autonomous manner, outside of the milieu of the mouse.

**Figure 6.**
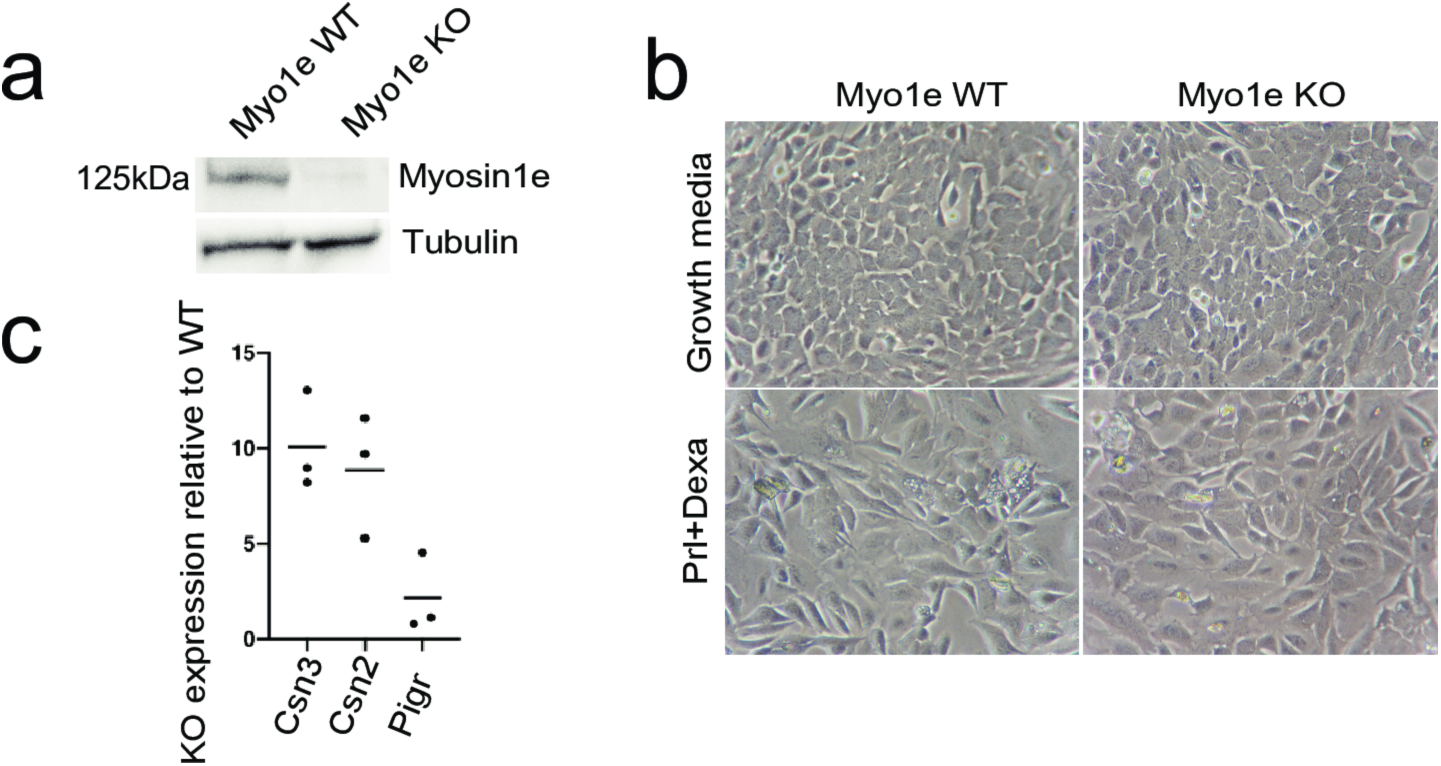
Py230 cells depleted of Myo1e have elevated milk-gene expression. (a) Western blot showing Myo1e expression in WT and Myo1e-KO Py230 cells. (b) Phase-contrast microscopy images of WT and Myo1e KO Py230 cells treated with growth media or growth media containing Prolactin (Prl) + Dexamethasone. (c) qRT-PCR results showing relative expression of Csn2, Csn3 and Pigr in KO Py230 cells treated with Prl + Dexamethasone compared to WT Py230 cells treated with Prl + Dexamethasone.

### Myo1e expression affects paracellular junctional strength of Py230 cells

Myo1e has been shown to regulate cell-cell contacts in kidney epithelial cells (48). Immunohistochemical staining of MMTV-PyMT tumors suggested that Myo1e localizes to E-cadherin-positive cell-cell junctions of tumor cells (3). We tested if Myo1e localizes to the cell-cell junction of Py230 cells by transient expression of mEmerald-tagged Myosin-1e in Myo1e-KO Py230 cells and found that mEmerald-tagged Myo1e localized to cell-cell contacts (Fig 7a). Gene-set enrichment analysis of RNA-seq data between Myo1e-WT and KO tumors indicated that cell-cell junction-associated genes were in the top 5 most enriched cellular compartments and biological processes affected by Myo1e expression (Table 1-2). We thus hypothesized that Myo1e expression influences cell-cell junction integrity in mammary tumor cells. We tested whether Myo1e expression affected cell-cell or cell-ECM junctions of mammary cancer cells using Electric-Cell-Substrate Impedance sensing (ECIS) (49, 50). As previously reported (49, 50), use of ECIS allows measurement of several parameters, including membrane capacitance, paracellular junctional strength and basal cell adhesion to substrate. Electrical current flow through the cell membrane was not statistically different between Myo1e-WT and KO Py230 cells (Fig 7b). This suggested that cell coverage over the electrodes as well as the cellular membrane properties were similar between Myo1e-WT and KO cells. Examining electrical current flow between the cell and substrate suggested that there was also no significant change in basal adhesion (Fig 7c). However, when current flow through cell-cell junctions was examined, there was a significant change in paracellular junctional strength between WT and KO Py230 cells (Fig 7d). Specifically, Myo1e-KO Py230 cells exhibited lower paracellular flow of the current through cell-cell junctions, which indicated stronger epithelial paracellular junctional strength compared to Myo1e-WT Py230 cells (Fig 7d). These findings suggest that Myo1e expression changes the properties of mammary tumor cell-cell junctions in a cell-autonomous manner.

**Figure 7.**
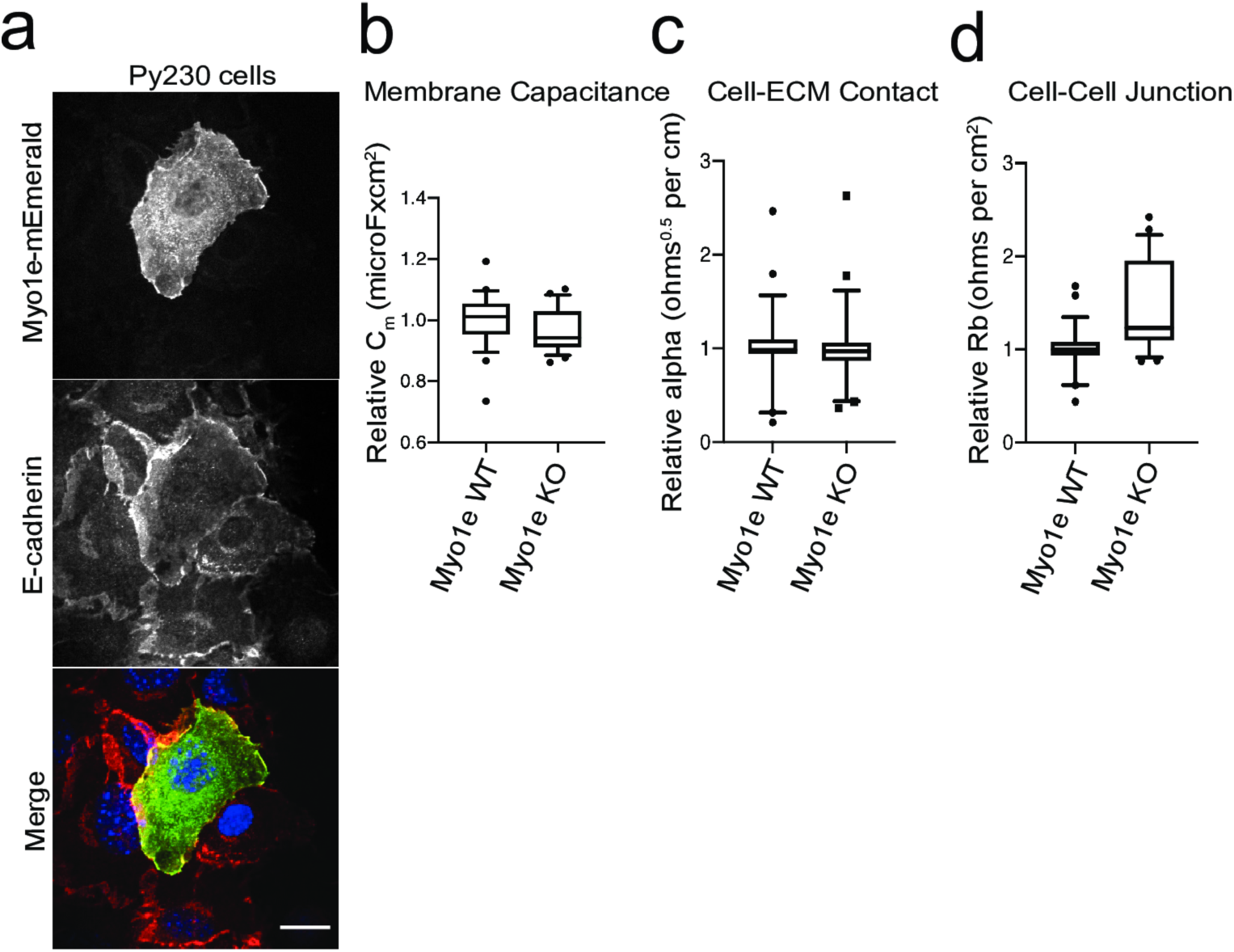
Myo1e expression affects paracellular junctional strength of Py230 cells. (a) Immunofluorescence staining for E-cadherin (red) in Py230 cells transiently expressing mEmerald-Myo1e (green). Nuclear DAPI staining is shown in blue. (b) ECIS measurement of membrane capacitance of WT and Myo1e-KO Py230 cells. (c) ECIS measurement of cell-ECM contact of WT and Myo1e-KO Py230 cells. (d) ECIS measurement of cell-cell junction barrier strength of WT and Myo1e-KO Py230 cells, p<0.001. Student’s T-test was used for statistical analysis. Scale bar: (a) 20µm.

## Discussion

In this paper, we aimed to identify specific molecular features of the mammary tumors that form in the MMTV-PyMT mouse model of tumorigenesis in wild-type and Myo1e-KO mice. We show that the expression of the unconventional myosin Myo1e is associated with a tumor transcriptional profile characterized by upregulation of tumor-promoting transcripts, and that the lack of Myo1e promotes mammary tumor cell differentiation and increases the strength of cell-cell junctions.

Transcriptomic profiling conducted in our study revealed that Myo1e expression in MMTV-PyMT tumors was associated with higher expression of genes linked to tumor growth and invasion compared to the tumors lacking Myo1e. Specifically, Tln2 and Myh10, which are upregulated in Myo1e-expressing tumors, have previously been shown to positively regulate cancer cell migration or invasion (51–54). The genes Ybx2, Arg2, Wnt4, Azgp1 and Nlgn3, which are significantly upregulated in Myo1e WT tumors, have been associated with tumor proliferation (55–59). Podxl, which is upregulated in Myo1e WT tumors, has an unclear role in cancer proliferation as Podxl expression has been associated with increased breast cancer cell growth and metastasis in xenograft models of breast cancer (60), but other studies have shown that it affects cell extravasation following EMT with no observed effect on cell growth (61). Conversely, the genes Rassf1, Rasl10a and Osgin1, which were all upregulated in the Myo1e-KO tumors, have been negatively associated with cancer cell proliferation and positively associated with promoting cell death (62–65). Dusp14 is also upregulated in the Myo1e-KO tumors and is negatively associated with the MAPK growth pathway (66). As a follow-up to these findings, we examined expression of proliferation markers Cyclin D1 and Ki-67 in tumors and tumor cells. Both Cyclin D1 distribution and Ki67 expression were indicative of a more proliferative state in the Myo1e-expressing tumors, in agreement with our earlier study (3) and with our RNA-seq findings. Primary tumor cells extracted from Myo1e-expressing mice had a higher percentage of cells with nuclear Cyclin D1 compared to Myo1e-KO primary tumor cells, suggesting that the changes in proliferation were also maintained outside of the tumor milieu. The mechanisms linking Myo1e expression with tumor cell proliferation remain to be elucidated, although we have observed that Myo1e-expressing primary tumor cells had higher nuclear enrichment of the transcriptional co-activator YAP compared to Myo1e-KO primary tumor cells, suggesting that Hippo/YAP signaling could mediate the effects of Myo1e. This observation is concordant with the findings that YAP expression affects both MMTV-PyMT tumor proliferation as well as the ability of mammary cells to terminally differentiate during pregnancy (34).

We have found that tumors depleted of Myo1e exhibit increased markers of differentiation compared to tumors expressing Myo1e. These markers include preserved mammary tissue architecture, expression of tumor-suppressive genes and expression and secretion of milk-associated proteins. Loss of epithelial differentiation frequently accompanies the development of malignant tumors (67, 68). In the MMTV-PyMT model, loss of expression of the epithelial cell markers E-cadherin and EpCAM, as well as upregulation of transcriptional profiles associated with EMT-mediating transcription factors (such as Slug/Snail/Zeb1) mark induction of EMT (69, 70). Since partial EMT or hybrid EMT can be observed in tumor cells (70), (71), interpreting EMT or neoplastic transformation as strictly associated with the loss of E-cadherin may represent an oversimplification, and additional markers of tumor aggressiveness may be predictive of patient outcomes. Interestingly, expression of lactation-associated proteins such as caseins and alpha-lactalbumin was shown to correlate with the increased epithelial differentiation of MMTV-PyMT tumor cells and a reduction in the metastatic potential (43–45, 72). The correlation between the expression of milk-associated proteins and lack of tumor aggressiveness in the MMTV-PyMT model further supports the idea that breastfeeding is protective against breast cancer via promoting differentiation of mammary cells (73). Our RNA-seq analysis showed that DEGs associated with extracellular (secreted) proteins were the most enriched cellular component among those affected by the presence or absence of Myo1e (Table 1). We found an accumulation of albumin, casein, and other secreted major milk-associated proteins in the Myo1e KO tumor cystic fluid. In addition, Myo1e KO mammary tumor cells produce an IHC signal from anti-milk staining that is not observed in less-differentiated Myo1e expressing tumors, supporting the results of the RNA-seq analysis of tumors and MS/MS analysis of KO cystic fluid. Moreover, Py230 cells depleted of Myo1e have increased expression of casein genes when treated with differentiation-inducing hormone (Prolactin), suggesting that Myo1e affects the expression of some milk-component genes in a cell-autonomous manner. These findings are in line with previous observations in showing that Myo1e expression is inversely associated with the differentiation state of intestinal epithelial cells, with higher expression of Myo1e in the intestinal crypts containing undifferentiated stem cells (15). Thus, overall, depletion of Myo1e may result in more differentiated state of epithelial cells and be protective against malignant transformation and EMT.

Interestingly, the effects of Myo1e depletion on tumor cell differentiation are similar to the effects of inhibiting Rankl signaling (45). Since Myo1e activity has been shown to be regulated downstream of Rankl signaling, with Rankl affecting both Myo1e expression and Myo1e-membrane association in osteoclasts (11), it is possible that the changes in differentiation observed in MMTV-PyMT tumor cells upon depletion of Rankl (45) may be mediated in part by the downstream changes to Myo1e expression and activity. Similarly to Myo1e KO, Rankl inhibition in the MMTV-PyMT model promotes tumor cell differentiation into apocrine, milk-producing cells, leading to a marked decrease in tumorigenicity (45). Rank/Rankl signaling is a well-studied and therapeutically targeted pathway that contributes to breast cancer initiation, progression, survival and metastasis (74, 75). Targeting Rankl via monoclonal antibody denosumab has recently been tested in phase-3 clinical trials for prevention of breast cancer recurrence and metastasis, with mixed results (76, 77). To better understand the effects of Rank signaling and attempt to identify a more targeted pathway for breast cancer treatment, future work should examine the relationship between Rank signaling and Myo1e activity in breast cancer cells.

Myo1e has been shown to modulate cell-cell contact formation in kidney epithelial cells (48), cell-extracellular matrix (ECM) adhesion (13, 78), and the stability of invadopodia (specialized adhesion structures) (79). Our transcriptomic analysis revealed that DEGs associated with cell-ECM adhesion, lipid transport and cell-cell adhesion were the top three enriched biological processes regulated by the presence or absence of Myo1e. In this study, we have also demonstrated that Myo1e influences cell-cell junction properties in a cell-autonomous manner by measuring paracellular junction strength in Py230 cells. Expression of the tight junction scaffolding protein ZO-1, which is a binding partner of Myo1e (48), was found to be lost in human breast cancer tissue (80, 81). Moreover, the expression of ZO-1 in human breast cancer tissue is found to significantly correlate with the differentiation state and progression of mammary tumors (80, 81). Similarly to human breast cancer, progressive loss of ZO-1 and cell polarity also occurs during tumor progression in the MMTV-PyMT mouse model of tumorigenesis (82) while maintenance of cell-cell junction integrity and activation of E-cadherin impedes MMTV-PyMT progression towards metastatic tumors (71). Cell-cell junctions affect both cancer progression and metastasis via being a signaling hub for pathways such as Hippo(YAP/TAZ), WNT, and Rac and by serving as structural components supporting cell polarity (83, 84). It is thus possible that Myo1e influences cell-cell junction stability or adhesion-regulated Hippo signaling and, as a result, lack of Myo1e may affect tumor progression.

In addition to changing cell-cell junction properties, Myo1e may be affecting pro-tumorigenic signaling via recruitment of pro-inflammatory immune cells (1, 12). We found decreased abundance of tumor associated intraepithelial macrophages in mice depleted of Myo1e expression when compared to tumors of WT mice. This is consistent with a study that found that the depletion of tumor associated macrophages in the MMTV-PyMT model of breast cancer is associated with reduction in tumor proliferation and metastasis through reduction in angiogenesis (36). Additionally, we found that the macrophage associated genes *Vsig4* and *Arg2* are differentially expressed between Myo1e WT and Myo1e KO tumors. Expression of *Vsig4*(39) and *Arg2*(85) has been identified to influence the M1/M2 activation state of macrophages. Thus, changes in the number of intra-epithelial macrophages or expression of M1/M2 activation-associated genes may therefore be a separate avenue via which Myo1e affects tumor progression.

Overall, our findings provide new insights into the effects of Myo1e loss on mammary tumor progression and suggest that Myo1e-mediated signaling may affect pathways involved in tumorigenesis and metastasis, including Hippo signaling, cell-cell adhesion remodeling, and macrophage infiltration. Some of these changes can be directly attributed to the effects of Myo1e loss on mammary tumor cells while others may result from an interplay between Myo1e-deficient tumors and Myo1e-deficient tumor microenvironment.

## Acknowledgments

We are thankful to Christopher Turner, Mariano Viapiano, and the members of the Sirotkin, Amack, and Pruyne labs for their helpful discussions regarding this project. We also thank Frank Middleton and Karen Gentile of the SUNY Molecular Analysis Core for their expert technical help with the RNA-seq experiments and analysis and for helpful discussion of the data. This project was supported by a NYS Health Dept. Peter T. Rowley Breast Cancer Scientific Research Project award, Carol M. Baldwin Breast Cancer Research Fund of Central New York Research Award, and Upstate Cancer Center pilot grant award to MK.

## Author Contributions

**Conceptualization:** Eric L Plante, Mira Krendel, Sharon Chase, Michael Garone.

**Data Curation:** Eric L Plante, Ebbing P de Jong, Theresa M Curtis.

**Formal Analysis:** Eric L Plante, Ebbing P de Jong, Theresa M Curtis.

**Investigation:** Eric L Plante, Sharon Chase.

**Methodology:** Eric L Plante, Mira Krendel, Sharon Chase.

**Supervision:** Mira Krendel.

**Writing – original draft:** Eric L Plante.

**Writing – review & editing:** Eric L Plante, Mira Krendel, Ebbing P de Jong, Theresa M Curtis, Sharon Chase, Michael Garone.

## Notes

### Competing Interest Statement

The authors have declared no competing interest.

## References

1. Ouderkirk JL, Krendel M. Non-muscle myosins in tumor progression, cancer cell invasion, and metastasis. Cytoskeleton (Hoboken). 2014;71(8):447–63.

2. Hallett RM, Dvorkin-Gheva A, Bane A, Hassell JA. A gene signature for predicting outcome in patients with basal-like breast cancer. Sci Rep. 2012;2:227.

3. Ouderkirk-Pecone JL, Goreczny GJ, Chase SE, Tatum AH, Turner CE, Krendel M. Myosin 1e promotes breast cancer malignancy by enhancing tumor cell proliferation and stimulating tumor cell de-differentiation. Oncotarget. 2016;7(29):46419–32.

4. Attalla S, Taifour T, Bui T, Muller W. Insights from transgenic mouse models of PyMT-induced breast cancer: recapitulating human breast cancer progression in vivo. Oncogene. 2021;40(3):475–91.

5. Davie SA, Maglione JE, Manner CK, Young D, Cardiff RD, MacLeod CL, et al. Effects of FVB/NJ and C57Bl/6J strain backgrounds on mammary tumor phenotype in inducible nitric oxide synthase deficient mice. Transgenic Res. 2007;16(2):193–201.

6. Pfefferle AD, Herschkowitz JI, Usary J, Harrell JC, Spike BT, Adams JR, et al. Transcriptomic classification of genetically engineered mouse models of breast cancer identifies human subtype counterparts. Genome Biol. 2013;14(11):R125.

7. Lin EY, Jones JG, Li P, Zhu L, Whitney KD, Muller WJ, et al. Progression to malignancy in the polyoma middle T oncoprotein mouse breast cancer model provides a reliable model for human diseases. Am J Pathol. 2003;163(5):2113–26.

8. Rennhack JP, To B, Swiatnicki M, Dulak C, Ogrodzinski MP, Zhang Y, et al. Integrated analyses of murine breast cancer models reveal critical parallels with human disease. Nat Commun. 2019;10(1):3261.

9. Dvornikov D, Schneider MA, Ohse S, Szczygiel M, Titkova I, Rosenblatt M, et al. Expression ratio of the TGFbeta-inducible gene MYO10 is prognostic for overall survival of squamous cell lung cancer patients and predicts chemotherapy response. Sci Rep. 2018;8(1):9517.

10. Tanimura S, Hashizume J, Arichika N, Watanabe K, Ohyama K, Takeda K, et al. ERK signaling promotes cell motility by inducing the localization of myosin 1E to lamellipodial tips. J Cell Biol. 2016;214(4):475–89.

11. Nakamura S, Masuyama R, Sakai K, Fukuda K, Takeda K, Tanimura S. SH3P2 suppresses osteoclast differentiation through restricting membrane localization of myosin 1E. Genes Cells. 2020;25(11):707–17.

12. Giron-Perez DA, Vadillo E, Schnoor M, Santos-Argumedo L. Myo1e modulates the recruitment of activated B cells to inguinal lymph nodes. J Cell Sci. 2020;133(5).

13. Heim JB, Squirewell EJ, Neu A, Zocher G, Sominidi-Damodaran S, Wyles SP, et al. Myosin-1E interacts with FAK proline-rich region 1 to induce fibronectin-type matrix. Proc Natl Acad Sci U S A. 2017;114(15):3933–8.

14. Krendel M, Kim SV, Willinger T, Wang T, Kashgarian M, Flavell RA, et al. Disruption of Myosin 1e promotes podocyte injury. J Am Soc Nephrol. 2009;20(1):86–94.

15. Skowron JF, Bement WM, Mooseker MS. Human brush border myosin-I and myosin-Ic expression in human intestine and Caco-2BBe cells. Cell Motil Cytoskeleton. 1998;41(4):308–24.

16. Wisniewski JR, Zougman A, Nagaraj N, Mann M. Universal sample preparation method for proteome analysis. Nat Methods. 2009;6(5):359–62.

17. Rappsilber J, Ishihama Y, Mann M. Stop and go extraction tips for matrix-assisted laser desorption/ionization, nanoelectrospray, and LC/MS sample pretreatment in proteomics. Anal Chem. 2003;75(3):663–70.

18. Romagnoli M, Bresson L, Di-Cicco A, Perez-Lanzon M, Legoix P, Baulande S, et al. Laminin-binding integrins are essential for the maintenance of functional mammary secretory epithelium in lactation. Development. 2020;147(4).

19. Rooney N, Wang P, Brennan K, Gilmore AP, Streuli CH. The Integrin-Mediated ILK-Parvin-alphaPix Signaling Axis Controls Differentiation in Mammary Epithelial Cells. J Cell Physiol. 2016;231(11):2408–17.

20. Liu K, Cheng L, Flesken-Nikitin A, Huang L, Nikitin AY, Pauli BU. Conditional knockout of fibronectin abrogates mouse mammary gland lobuloalveolar differentiation. Dev Biol. 2010;346(1):11–24.

21. Muschler J, Streuli CH. Cell-matrix interactions in mammary gland development and breast cancer. Cold Spring Harb Perspect Biol. 2010;2(10):a003202.

22. Bridgewater RE, Streuli CH, Caswell PT. Extracellular matrix promotes clathrin-dependent endocytosis of prolactin and STAT5 activation in differentiating mammary epithelial cells. Sci Rep. 2017;7(1):4572.

23. Akhtar N, Marlow R, Lambert E, Schatzmann F, Lowe ET, Cheung J, et al. Molecular dissection of integrin signalling proteins in the control of mammary epithelial development and differentiation. Development. 2009;136(6):1019–27.

24. Joyce JA, Pollard JW. Microenvironmental regulation of metastasis. Nat Rev Cancer. 2009;9(4):239–52.

25. Cai Y, Nogales-Cadenas R, Zhang Q, Lin JR, Zhang W, O’Brien K, et al. Transcriptomic dynamics of breast cancer progression in the MMTV-PyMT mouse model. BMC Genomics. 2017;18(1):185.

26. Patel OV, Casey T, Dover H, Plaut K. Homeorhetic adaptation to lactation: comparative transcriptome analysis of mammary, liver, and adipose tissue during the transition from pregnancy to lactation in rats. Funct Integr Genomics. 2011;11(1):193–202.

27. Huynh H, Ng CY, Ong CK, Lim KB, Chan TW. Cloning and characterization of a novel pregnancy-induced growth inhibitor in mammary gland. Endocrinology. 2001;142(8):3607–15.

28. Rosato R, Jammes H, Belair L, Puissant C, Kraehenbuhl JP, Djiane J. Polymeric-Ig receptor gene expression in rabbit mammary gland during pregnancy and lactation: evolution and hormonal regulation. Mol Cell Endocrinol. 1995;110(1-2):81–7.

29. Rincheval-Arnold A, Belair L, Djiane J. Developmental expression of pIgR gene in sheep mammary gland and hormonal regulation. J Dairy Res. 2002;69(1):13–26.

30. Boumahrou N, Chevaleyre C, Berri M, Martin P, Bellier S, Salmon H. An increase in milk IgA correlates with both pIgR expression and IgA plasma cell accumulation in the lactating mammary gland of PRM/Alf mice. J Reprod Immunol. 2012;96(1-2):25–33.

31. Zhou G, Soufan O, Ewald J, Hancock REW, Basu N, Xia J. NetworkAnalyst 3.0: a visual analytics platform for comprehensive gene expression profiling and meta-analysis. Nucleic Acids Res. 2019;47(W1):W234–W41.

32. Calses PC, Crawford JJ, Lill JR, Dey A. Hippo Pathway in Cancer: Aberrant Regulation and Therapeutic Opportunities. Trends Cancer. 2019;5(5):297–307.

33. van der Weyden L, Papaspyropoulos A, Poulogiannis G, Rust AG, Rashid M, Adams DJ, et al. Loss of RASSF1A synergizes with deregulated RUNX2 signaling in tumorigenesis. Cancer Res. 2012;72(15):3817–27.

34. Chen Q, Zhang N, Gray RS, Li H, Ewald AJ, Zahnow CA, et al. A temporal requirement for Hippo signaling in mammary gland differentiation, growth, and tumorigenesis. Genes Dev. 2014;28(5):432–7.

35. Nishio M, Otsubo K, Maehama T, Mimori K, Suzuki A. Capturing the mammalian Hippo: elucidating its role in cancer. Cancer Sci. 2013;104(10):1271–7.

36. Rumney RMH, Coffelt SB, Neale TA, Dhayade S, Tozer GM, Miller G. PyMT-Maclow: A novel, inducible, murine model for determining the role of CD68 positive cells in breast tumor development. PLoS One. 2017;12(12):e0188591.

37. Gil-Varea E, Urcelay E, Vilarino-Guell C, Costa C, Midaglia L, Matesanz F, et al. Exome sequencing study in patients with multiple sclerosis reveals variants associated with disease course. J Neuroinflammation. 2018;15(1):265.

38. Al-Mutairi MS, Cadalbert LC, McGachy HA, Shweash M, Schroeder J, Kurnik M, et al. MAP kinase phosphatase-2 plays a critical role in response to infection by Leishmania mexicana. PLoS Pathog. 2010;6(11):e1001192.

39. Li J, Diao B, Guo S, Huang X, Yang C, Feng Z, et al. VSIG4 inhibits proinflammatory macrophage activation by reprogramming mitochondrial pyruvate metabolism. Nat Commun. 2017;8(1):1322.

40. Lewis ND, Asim M, Barry DP, de Sablet T, Singh K, Piazuelo MB, et al. Immune evasion by Helicobacter pylori is mediated by induction of macrophage arginase II. J Immunol. 2011;186(6):3632–41.

41. Wenzel J, Ouderkirk JL, Krendel M, Lang R. Class I myosin Myo1e regulates TLR4-triggered macrophage spreading, chemokine release, and antigen presentation via MHC class II. Eur J Immunol. 2015;45(1):225–37.

42. Barger SR, James ML, Pellenz CD, Krendel M, Sirotkin V. Human myosin 1e tail but not motor domain replaces fission yeast Myo1 domains to support myosin-I function during endocytosis. Exp Cell Res. 2019;384(2):111625.

43. Arun G, Diermeier S, Akerman M, Chang KC, Wilkinson JE, Hearn S, et al. Differentiation of mammary tumors and reduction in metastasis upon Malat1 lncRNA loss. Genes Dev. 2016;30(1):34–51.

44. Hebbard L, Cecena G, Golas J, Sawada J, Ellies LG, Charbono A, et al. Control of mammary tumor differentiation by SKI-606 (bosutinib). Oncogene. 2011;30(3):301–12.

45. Yoldi G, Pellegrini P, Trinidad EM, Cordero A, Gomez-Miragaya J, Serra-Musach J, et al. RANK Signaling Blockade Reduces Breast Cancer Recurrence by Inducing Tumor Cell Differentiation. Cancer Res. 2016;76(19):5857–69.

46. Gyorffy B, Lanczky A, Eklund AC, Denkert C, Budczies J, Li Q, et al. An online survival analysis tool to rapidly assess the effect of 22,277 genes on breast cancer prognosis using microarray data of 1,809 patients. Breast Cancer Res Treat. 2010;123(3):725–31.

47. Bao L, Cardiff RD, Steinbach P, Messer KS, Ellies LG. Multipotent luminal mammary cancer stem cells model tumor heterogeneity. Breast Cancer Res. 2015;17(1):137.

48. Bi J, Chase SE, Pellenz CD, Kurihara H, Fanning AS, Krendel M. Myosin 1e is a component of the glomerular slit diaphragm complex that regulates actin reorganization during cell-cell contact formation in podocytes. Am J Physiol Renal Physiol. 2013;305(4):F532–44.

49. Robilliard LD, Kho DT, Johnson RH, Anchan A, O’Carroll SJ, Graham ES. The Importance of Multifrequency Impedance Sensing of Endothelial Barrier Formation Using ECIS Technology for the Generation of a Strong and Durable Paracellular Barrier. Biosensors (Basel). 2018;8(3).

50. Giaever I, Keese CR. Micromotion of mammalian cells measured electrically. Proc Natl Acad Sci U S A. 1991;88(17):7896–900.

51. Betapudi V. Myosin II motor proteins with different functions determine the fate of lamellipodia extension during cell spreading. PLoS One. 2010;5(1):e8560.

52. Vicente-Manzanares M, Ma X, Adelstein RS, Horwitz AR. Non-muscle myosin II takes centre stage in cell adhesion and migration. Nat Rev Mol Cell Biol. 2009;10(11):778–90.

53. Li L, Li X, Qi L, Rychahou P, Jafari N, Huang C. The role of talin2 in breast cancer tumorigenesis and metastasis. Oncotarget. 2017;8(63):106876–87.

54. Thomas DG, Yenepalli A, Denais CM, Rape A, Beach JR, Wang YL, et al. Non-muscle myosin IIB is critical for nuclear translocation during 3D invasion. J Cell Biol. 2015;210(4):583–94.

55. Niu X, Yang B, Liu F, Fang Q. LncRNA HOXA11-AS promotes OSCC progression by sponging miR-98-5p to upregulate YBX2 expression. Biomed Pharmacother. 2020;121:109623.

56. Sikora MJ, Jacobsen BM, Levine K, Chen J, Davidson NE, Lee AV, et al. WNT4 mediates estrogen receptor signaling and endocrine resistance in invasive lobular carcinoma cell lines. Breast Cancer Res. 2016;18(1):92.

57. Ji M, Li W, He G, Zhu D, Lv S, Tang W, et al. Zinc-alpha2-glycoprotein 1 promotes EMT in colorectal cancer by filamin A mediated focal adhesion pathway. J Cancer. 2019;10(22):5557–66.

58. Li Z, Gao W, Fei Y, Gao P, Xie Q, Xie J, et al. NLGN3 promotes neuroblastoma cell proliferation and growth through activating PI3K/AKT pathway. Eur J Pharmacol. 2019;857:172423.

59. Costa H, Xu X, Overbeek G, Vasaikar S, Patro CP, Kostopoulou ON, et al. Human cytomegalovirus may promote tumour progression by upregulating arginase-2. Oncotarget. 2016;7(30):47221–31.

60. Snyder KA, Hughes MR, Hedberg B, Brandon J, Hernaez DC, Bergqvist P, et al. Podocalyxin enhances breast tumor growth and metastasis and is a target for monoclonal antibody therapy. Breast Cancer Res. 2015;17:46.

61. Frose J, Chen MB, Hebron KE, Reinhardt F, Hajal C, Zijlstra A, et al. Epithelial-Mesenchymal Transition Induces Podocalyxin to Promote Extravasation via Ezrin Signaling. Cell Rep. 2018;24(4):962–72.

62. Ram RR, Mendiratta S, Bodemann BO, Torres MJ, Eskiocak U, White MA. RASSF1A inactivation unleashes a tumor suppressor/oncogene cascade with context-dependent consequences on cell cycle progression. Mol Cell Biol. 2014;34(12):2350–8.

63. Donninger H, Vos MD, Clark GJ. The RASSF1A tumor suppressor. J Cell Sci. 2007;120(Pt 18):3163–72.

64. Tsai CH, Shen YC, Chen HW, Liu KL, Chang JW, Chen PY, et al. Docosahexaenoic acid increases the expression of oxidative stress-induced growth inhibitor 1 through the PI3K/Akt/Nrf2 signaling pathway in breast cancer cells. Food Chem Toxicol. 2017;108(Pt A):276–88.

65. Elam C, Hesson L, Vos MD, Eckfeld K, Ellis CA, Bell A, et al. RRP22 is a farnesylated, nucleolar, Ras-related protein with tumor suppressor potential. Cancer Res. 2005;65(8):3117–25.

66. Li CY, Zhou Q, Yang LC, Chen YH, Hou JW, Guo K, et al. Dual-specificity phosphatase 14 protects the heart from aortic banding-induced cardiac hypertrophy and dysfunction through inactivation of TAK1-P38MAPK/-JNK1/2 signaling pathway. Basic Res Cardiol. 2016;111(2):19.

67. Pattabiraman DR, Weinberg RA. Targeting the Epithelial-to-Mesenchymal Transition: The Case for Differentiation-Based Therapy. Cold Spring Harb Symp Quant Biol. 2016;81:11–9.

68. Bhatia S, Wang P, Toh A, Thompson EW. New Insights Into the Role of Phenotypic Plasticity and EMT in Driving Cancer Progression. Front Mol Biosci. 2020;7:71.

69. Ye X, Tam WL, Shibue T, Kaygusuz Y, Reinhardt F, Ng Eaton E, et al. Distinct EMT programs control normal mammary stem cells and tumour-initiating cells. Nature. 2015;525(7568):256–60.

70. Pastushenko I, Brisebarre A, Sifrim A, Fioramonti M, Revenco T, Boumahdi S, et al. Identification of the tumour transition states occurring during EMT. Nature. 2018;556(7702):463–8.

71. Na TY, Schecterson L, Mendonsa AM, Gumbiner BM. The functional activity of E-cadherin controls tumor cell metastasis at multiple steps. Proc Natl Acad Sci U S A. 2020;117(11):5931–7.

72. Kouros-Mehr H, Bechis SK, Slorach EM, Littlepage LE, Egeblad M, Ewald AJ, et al. GATA-3 links tumor differentiation and dissemination in a luminal breast cancer model. Cancer Cell. 2008;13(2):141–52.

73. Kotsopoulos J, Lubinski J, Salmena L, Lynch HT, Kim-Sing C, Foulkes WD, et al. Breastfeeding and the risk of breast cancer in BRCA1 and BRCA2 mutation carriers. Breast Cancer Res. 2012;14(2):R42.

74. Wu X, Li F, Dang L, Liang C, Lu A, Zhang G. RANKL/RANK System-Based Mechanism for Breast Cancer Bone Metastasis and Related Therapeutic Strategies. Front Cell Dev Biol. 2020;8:76.

75. Palafox M, Ferrer I, Pellegrini P, Vila S, Hernandez-Ortega S, Urruticoechea A, et al. RANK induces epithelial-mesenchymal transition and stemness in human mammary epithelial cells and promotes tumorigenesis and metastasis. Cancer Res. 2012;72(11):2879–88.

76. Gnant M, Pfeiler G, Steger GG, Egle D, Greil R, Fitzal F, et al. Adjuvant denosumab in postmenopausal patients with hormone receptor-positive breast cancer (ABCSG-18): disease-free survival results from a randomised, double-blind, placebo-controlled, phase 3 trial. Lancet Oncol. 2019;20(3):339–51.

77. Coleman R, Finkelstein DM, Barrios C, Martin M, Iwata H, Hegg R, et al. Adjuvant denosumab in early breast cancer (D-CARE): an international, multicentre, randomised, controlled, phase 3 trial. Lancet Oncol. 2020;21(1):60–72.

78. Gupta P, Gauthier NC, Cheng-Han Y, Zuanning Y, Pontes B, Ohmstede M, et al. Myosin 1E localizes to actin polymerization sites in lamellipodia, affecting actin dynamics and adhesion formation. Biol Open. 2013;2(12):1288–99.

79. Ouderkirk JL, Krendel M. Myosin 1e is a component of the invadosome core that contributes to regulation of invadosome dynamics. Exp Cell Res. 2014;322(2):265–76.

80. Hoover KB, Liao SY, Bryant PJ. Loss of the tight junction MAGUK ZO-1 in breast cancer: relationship to glandular differentiation and loss of heterozygosity. Am J Pathol. 1998;153(6):1767–73.

81. Martin TA, Watkins G, Mansel RE, Jiang WG. Loss of tight junction plaque molecules in breast cancer tissues is associated with a poor prognosis in patients with breast cancer. Eur J Cancer. 2004;40(18):2717–25.

82. Halaoui R, Rejon C, Chatterjee SJ, Szymborski J, Meterissian S, Muller WJ, et al. Progressive polarity loss and luminal collapse disrupt tissue organization in carcinoma. Genes Dev. 2017;31(15):1573–87.

83. Knights AJ, Funnell AP, Crossley M, Pearson RC. Holding Tight: Cell Junctions and Cancer Spread. Trends Cancer Res. 2012;8:61–9.

84. Garcia MA, Nelson WJ, Chavez N. Cell-Cell Junctions Organize Structural and Signaling Networks. Cold Spring Harb Perspect Biol. 2018;10(4).

85. Ming XF, Rajapakse AG, Yepuri G, Xiong Y, Carvas JM, Ruffieux J, et al. Arginase II Promotes Macrophage Inflammatory Responses Through Mitochondrial Reactive Oxygen Species, Contributing to Insulin Resistance and Atherogenesis. J Am Heart Assoc. 2012;1(4):e000992.

