## Supplemental data for "Loss of Myosin-1e biases MMTV-PyMT induced breast cancer towards a differentiated and secretory state"

a

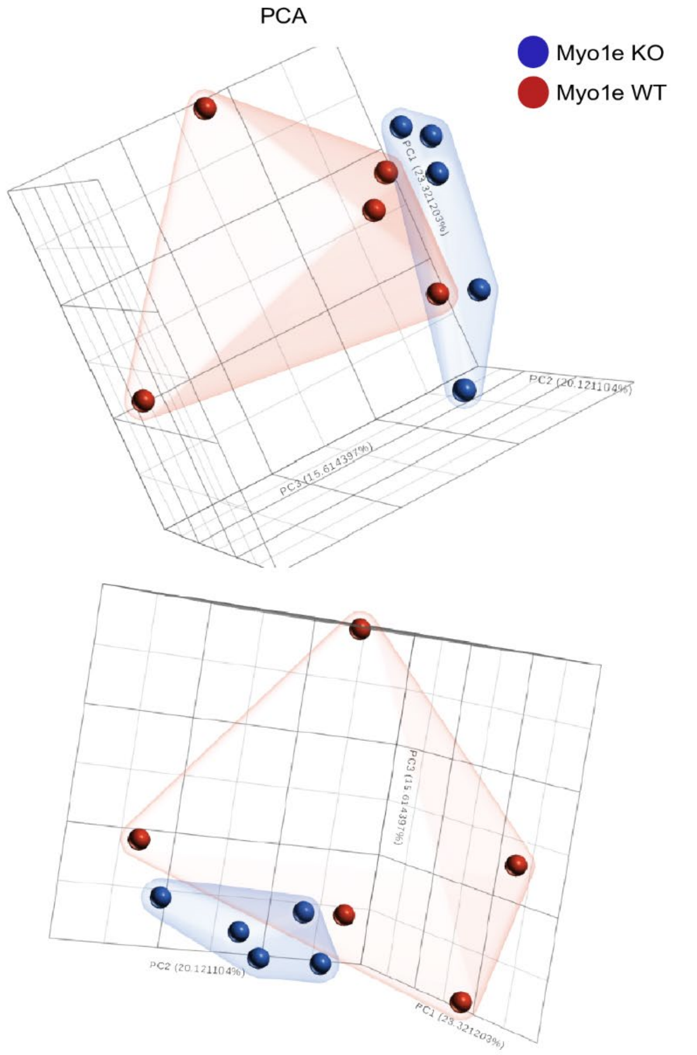
PCA
Myo1e KO
Myo1e WT
PC1 (23.321203%)
PC2 (20.121104%)
PC3 (15.614397%)
PC3 (15.614397%)
PC2 (20.121104%)
PC1 (23.321203%)

b

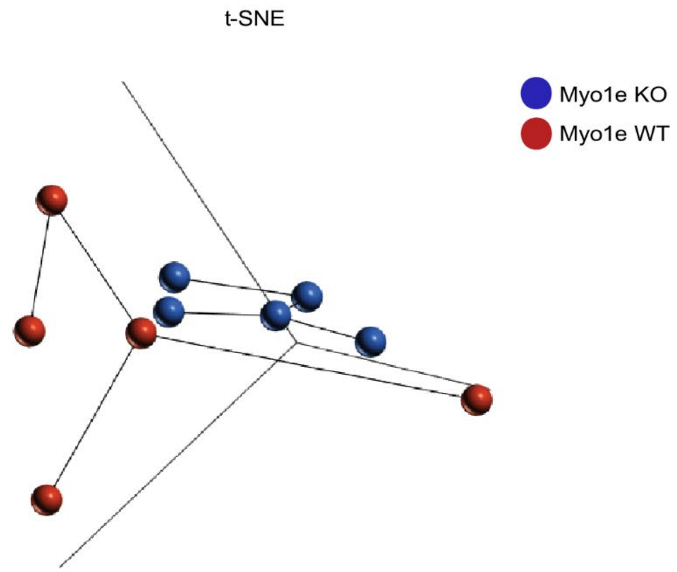
t-SNE
Myo1e KO
Myo1e WT

Supplemental Figure 1) **Statistical analysis of differentially expressed genes.** (a) Principal components analysis results suggest a distinct separation between differentially expressed genes in Myo1e WT and KO tumor tissue. (b) T-distributed stochastic neighbor embedding (t-SNE) analysis results also suggest a distinct separation between differentially expressed genes in Myo1e WT and KO tumor tissue. WT data points are shown in red and Myo1e KO data points are shown in blue.

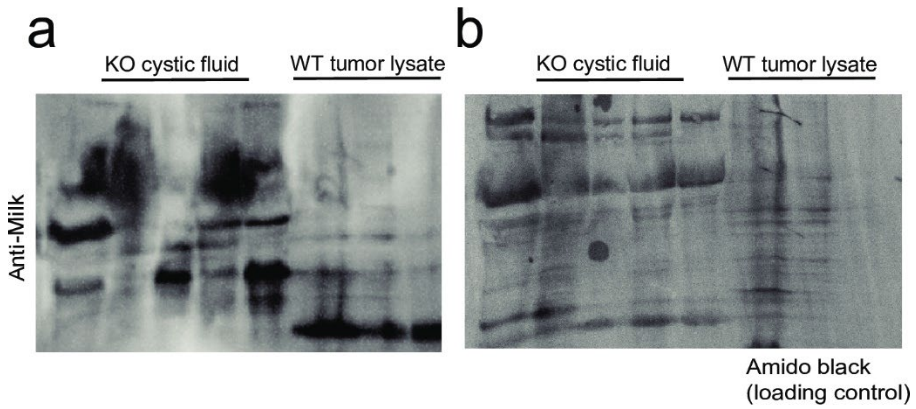


Supplemental Figure 2) **Western blot analysis of milk-associated proteins.** (a) Western blot analysis of tumor lysates from three Myo1e WT tumors and tumor cystic fluid from five Myo1e KO mice using antibodies against mouse milk-associated proteins. (b) Amido black staining to visualize total protein loading of samples shown in a.

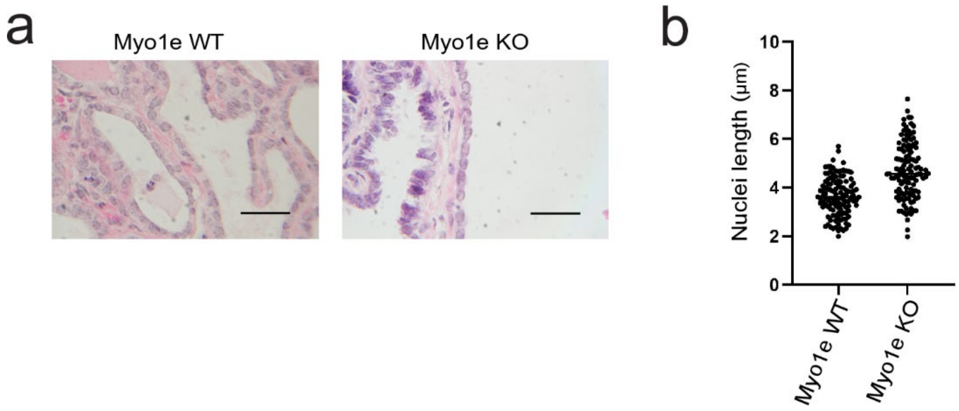


Supplemental Figure 3) **Nuclear length measurements of tumor cells.** (a) Images of epithelial tumor cells from H&E-stained FFPE tumor tissue from Myo1e WT tumors (left) and Myo1e KO tumors (right). (b) Measurement of nuclear length along the apical-basal axis. Each data points represents a measurement from an individual tumor cell. Scale bar: (a) 25µm.

| Gene ID | Chromosome | Start | Stop | Strand | Total counts | Average coverage | Q-value (KO vs. WT) | P-value (KO vs. WT) | FDR step up (KO vs. WT) | Ratio (KO vs. WT) | Fold change (KO vs. WT) | LSMean(KO) (KO vs. WT) | LSMean(WT) (KO vs. WT) | FC 95% lower limit (KO vs. WT) | FC 95% upper limit (KO vs. WT) |
|---|---|---|---|---|---|---|---|---|---|---|---|---|---|---|---|
| 1700030C10Rik | 12 | 20804392 | 20815780 | - | 3.55E+00 | 2.10E-01 | 1.56E-02 | 0.000001 | 1.67E-02 | 8.75E+00 | 8.75E+00 | 6.38E-01 | 7.29E-02 | 5.97E+00 | 1.28E+01 |
| Chit1 | 1 | 1.34E+08 | 1.34E+08 | + | 4.88E+00 | 2.51E-01 | 7.19E-01 | 0.003431 | 7.73E-01 | 7.08E+00 | 7.08E+00 | 8.54E-01 | 1.21E-01 | 2.12E+00 | 2.37E+01 |
| Vsig4 | X | 96247203 | 96293439 | - | 6.88E-02 | 4.20E-03 | 7.87E-01 | 0.007499 | 8.45E-01 | 5.13E+00 | 5.13E+00 | 1.15E-02 | 2.24E-03 | 1.97E+00 | 1.34E+01 |
| Fcna | 2 | 25624667 | 25627975 | - | 4.49E+00 | 2.60E-01 | 7.87E-01 | 0.007962 | 8.45E-01 | 5.10E+00 | 5.10E+00 | 7.50E-01 | 1.47E-01 | 1.57E+00 | 1.65E+01 |
| Gm16325 | 3 | 1.47E+08 | 1.47E+08 | + | 4.22E-01 | 2.77E-02 | 7.19E-01 | 0.003523 | 7.73E-01 | 4.79E+00 | 4.79E+00 | 6.98E-02 | 1.46E-02 | 2.00E+00 | 1.15E+01 |
| Lalba | 15 | 98480400 | 98482684 | - | 3.05E+03 | 2.10E+02 | 7.87E-01 | 0.008289 | 8.45E-01 | 3.47E+00 | 3.47E+00 | 4.73E+02 | 1.36E+02 | 1.52E+00 | 7.93E+00 |
| Rasl10a | 11 | 5058128 | 5060384 | + | 2.37E+00 | 1.89E-01 | 7.19E-01 | 0.003169 | 7.73E-01 | 3.27E+00 | 3.27E+00 | 3.62E-01 | 1.11E-01 | 1.74E+00 | 6.12E+00 |
| Apol7a | 15 | 77388217 | 77399111 | - | 4.40E+01 | 3.57E+00 | 7.87E-01 | 0.007732 | 8.45E-01 | 2.98E+00 | 2.98E+00 | 6.59E+00 | 2.22E+00 | 1.47E+00 | 6.03E+00 |
| Apol7c | 15 | 77524867 | 77533316 | - | 1.11E+00 | 9.66E-02 | 7.48E-01 | 0.004258 | 8.04E-01 | 2.36E+00 | 2.36E+00 | 1.55E-01 | 6.59E-02 | 1.43E+00 | 3.89E+00 |
| Snora34 | 15 | 98519717 | 98519835 | - | 3.98E+01 | 3.58E+00 | 7.19E-01 | 0.002739 | 7.73E-01 | 2.22E+00 | 2.22E+00 | 5.49E+00 | 2.47E+00 | 1.44E+00 | 3.44E+00 |
| Sept5 | 16 | 18621811 | 18629939 | - | 2.62E+01 | 2.40E+00 | 7.84E-01 | 0.005505 | 8.43E-01 | 1.98E+00 | 1.98E+00 | 3.49E+00 | 1.76E+00 | 1.32E+00 | 2.99E+00 |
| Pigr | 1 | 1.31E+08 | 1.31E+08 | + | 1.13E+02 | 1.04E+01 | 7.87E-01 | 0.007923 | 8.45E-01 | 1.98E+00 | 1.98E+00 | 1.51E+01 | 7.61E+00 | 1.26E+00 | 3.10E+00 |
| Gm12915 | 4 | 1.25E+08 | 1.25E+08 | - | 8.32E-01 | 7.74E-02 | 7.87E-01 | 0.008155 | 8.45E-01 | 1.85E+00 | 1.85E+00 | 1.08E-01 | 5.84E-02 | 1.25E+00 | 2.73E+00 |
| Gm6548 | 17 | 78850505 | 78852545 | - | 1.33E+03 | 1.27E+02 | 1.69E-02 | 0.000003 | 1.82E-02 | 1.85E+00 | 1.85E+00 | 1.73E+02 | 9.35E+01 | 1.64E+00 | 2.09E+00 |
| Apol7b | 15 | 77422209 | 77447461 | - | 5.27E-01 | 4.95E-02 | 7.19E-01 | 0.001716 | 7.73E-01 | 1.83E+00 | 1.83E+00 | 6.81E-02 | 3.72E-02 | 1.35E+00 | 2.48E+00 |
| Apol7e | 15 | 77698889 | 77719289 | + | 7.94E-01 | 7.43E-02 | 7.87E-01 | 0.006934 | 8.45E-01 | 1.78E+00 | 1.78E+00 | 1.02E-01 | 5.70E-02 | 1.23E+00 | 2.60E+00 |
| Osgin1 | 8 | 1.19E+08 | 1.19E+08 | + | 2.17E+01 | 2.04E+00 | 7.87E-01 | 0.005918 | 8.45E-01 | 1.78E+00 | 1.78E+00 | 2.78E+00 | 1.56E+00 | 1.25E+00 | 2.54E+00 |
| 1110008P14Rik | 2 | 32379101 | 32381916 | - | 4.89E+02 | 4.71E+01 | 7.19E-01 | 0.001711 | 7.73E-01 | 1.58E+00 | 1.58E+00 | 5.98E+01 | 3.79E+01 | 1.26E+00 | 1.98E+00 |
| Rassf1 | 9 | 1.08E+08 | 1.08E+08 | + | 5.75E+01 | 5.57E+00 | 7.19E-01 | 0.002370 | 7.73E-01 | 1.55E+00 | 1.55E+00 | 6.99E+00 | 4.52E+00 | 1.23E+00 | 1.94E+00 |
| Phactr1 | 13 | 42680799 | 43138513 | + | 1.11E+00 | 1.08E-01 | 7.87E-01 | 0.007852 | 8.45E-01 | 1.51E+00 | 1.51E+00 | 1.34E-01 | 8.88E-02 | 1.16E+00 | 1.96E+00 |
| Dusp14 | 11 | 84048045 | 84068358 | - | 4.67E+00 | 4.53E-01 | 7.19E-01 | 0.003108 | 7.73E-01 | 1.51E+00 | 1.51E+00 | 5.61E-01 | 3.73E-01 | 1.20E+00 | 1.89E+00 |
| Zfp850 | 7 | 27984471 | 28014116 | - | 2.37E+00 | 2.28E-01 | 7.87E-01 | 0.008270 | 8.45E-01 | 6.52E-01 | -1.53E+00 | 1.87E-01 | 2.87E-01 | -2.04E+00 | -1.15E+00 |
| Pigl | 11 | 62458460 | 62513901 | + | 1.15E+00 | 1.12E-01 | 7.19E-01 | 0.003060 | 7.73E-01 | 6.47E-01 | -1.55E+00 | 9.06E-02 | 1.40E-01 | -1.97E+00 | -1.22E+00 |
| Rsad1 | 11 | 94539798 | 94549208 | - | 1.28E+01 | 1.24E+00 | 6.75E-01 | 0.000587 | 7.25E-01 | 6.46E-01 | -1.55E+00 | 1.01E+00 | 1.56E+00 | -1.86E+00 | -1.29E+00 |
| Abhd14a | 9 | 1.06E+08 | 1.06E+08 | - | 2.81E+01 | 2.70E+00 | 7.87E-01 | 0.008340 | 8.45E-01 | 6.39E-01 | -1.56E+00 | 2.19E+00 | 3.43E+00 | -2.10E+00 | -1.16E+00 |
| Myzap | 9 | 71504347 | 71592361 | - | 1.46E+00 | 1.38E-01 | 7.33E-01 | 0.004120 | 7.87E-01 | 5.78E-01 | -1.73E+00 | 1.07E-01 | 1.85E-01 | -2.38E+00 | -1.25E+00 |
| Mycbpap | 11 | 94501347 | 94521503 | - | 3.82E+00 | 3.61E-01 | 7.67E-01 | 0.004463 | 8.24E-01 | 5.76E-01 | -1.73E+00 | 2.79E-01 | 4.85E-01 | -2.41E+00 | -1.25E+00 |
| Ccdc155 | 7 | 45183676 | 45204893 | - | 2.30E-01 | 2.15E-02 | 7.49E-01 | 0.004312 | 8.05E-01 | 5.69E-01 | -1.76E+00 | 1.67E-02 | 2.93E-02 | -2.45E+00 | -1.26E+00 |
| Chd3os | 11 | 69340769 | 69342648 | + | 8.97E+00 | 8.49E-01 | 7.19E-01 | 0.001981 | 7.73E-01 | 5.60E-01 | -1.78E+00 | 6.44E-01 | 1.15E+00 | -2.38E+00 | -1.34E+00 |
| Tmem2 | 19 | 21778340 | 21858361 | + | 1.12E+01 | 1.04E+00 | 8.23E-01 | 0.009644 | 8.84E-01 | 5.55E-01 | -1.80E+00 | 7.97E-01 | 1.44E+00 | -2.67E+00 | -1.21E+00 |
| Mks1 | 11 | 87853225 | 87863680 | + | 8.14E+00 | 7.56E-01 | 7.87E-01 | 0.008352 | 8.45E-01 | 5.52E-01 | -1.81E+00 | 5.79E-01 | 1.05E+00 | -2.70E+00 | -1.21E+00 |
| Tln2 | 9 | 67217085 | 67559704 | - | 3.58E+00 | 3.34E-01 | 7.92E-01 | 0.008609 | 8.51E-01 | 5.50E-01 | -1.82E+00 | 2.54E-01 | 4.62E-01 | -2.70E+00 | -1.22E+00 |
| Podxl | 6 | 31519493 | 31563938 | - | 2.28E+01 | 2.13E+00 | 7.87E-01 | 0.006819 | 8.45E-01 | 5.48E-01 | -1.82E+00 | 1.62E+00 | 2.95E+00 | -2.63E+00 | -1.26E+00 |
| Myh10 | 11 | 68691915 | 68816625 | + | 4.61E+00 | 4.27E-01 | 8.23E-01 | 0.009874 | 8.84E-01 | 5.27E-01 | -1.90E+00 | 3.18E-01 | 6.03E-01 | -2.88E+00 | -1.25E+00 |
| Map10 | 8 | 1.26E+08 | 1.26E+08 | + | 3.47E+00 | 3.25E-01 | 6.75E-01 | 0.000450 | 7.25E-01 | 5.20E-01 | -1.92E+00 | 2.37E-01 | 4.57E-01 | -2.51E+00 | -1.47E+00 |
| Psmc3ip | 11 | 1.01E+08 | 1.01E+08 | - | 1.64E+01 | 1.53E+00 | 7.19E-01 | 0.001194 | 7.73E-01 | 5.18E-01 | -1.93E+00 | 1.12E+00 | 2.16E+00 | -2.61E+00 | -1.43E+00 |
| Gimap9 | 6 | 48676135 | 48678705 | + | 1.91E+01 | 1.75E+00 | 7.79E-01 | 0.005058 | 8.37E-01 | 5.14E-01 | -1.95E+00 | 1.29E+00 | 2.52E+00 | -2.93E+00 | -1.30E+00 |
| Nr2f2 | 7 | 70351950 | 70366747 | - | 4.85E+00 | 4.49E-01 | 6.75E-01 | 0.000548 | 7.25E-01 | 5.09E-01 | -1.96E+00 | 3.27E-01 | 6.42E-01 | -2.61E+00 | -1.48E+00 |
| Steap1 | 5 | 5736322 | 5749318 | - | 1.86E+00 | 1.71E-01 | 7.33E-01 | 0.004099 | 7.87E-01 | 5.08E-01 | -1.97E+00 | 1.25E-01 | 2.47E-01 | -2.94E+00 | -1.32E+00 |
| Mrgpre | 7 | 1.44E+08 | 1.44E+08 | - | 2.97E+00 | 2.69E-01 | 8.23E-01 | 0.009709 | 8.84E-01 | 4.99E-01 | -2.01E+00 | 1.98E-01 | 3.97E-01 | -3.20E+00 | -1.26E+00 |
| Ybx2 | 11 | 69935899 | 69941602 | + | 4.02E+00 | 3.63E-01 | 7.87E-01 | 0.007720 | 8.45E-01 | 4.89E-01 | -2.04E+00 | 2.64E-01 | 5.39E-01 | -3.23E+00 | -1.29E+00 |
| Pxylp1 | 9 | 96823343 | 96889475 | - | 3.68E+00 | 3.29E-01 | 7.92E-01 | 0.008605 | 8.51E-01 | 4.71E-01 | -2.12E+00 | 2.36E-01 | 5.00E-01 | -3.44E+00 | -1.31E+00 |
| B3gnt7 | 1 | 86303221 | 86307306 | + | 8.99E+00 | 8.09E-01 | 7.19E-01 | 0.001216 | 7.73E-01 | 4.42E-01 | -2.26E+00 | 5.52E-01 | 1.25E+00 | -3.31E+00 | -1.54E+00 |
| Xk | X | 9272784 | 9313246 | + | 1.44E+00 | 1.25E-01 | 7.87E-01 | 0.005801 | 8.45E-01 | 4.19E-01 | -2.39E+00 | 8.52E-02 | 2.03E-01 | -4.12E+00 | -1.39E+00 |
| Lpcat4 | 2 | 1.12E+08 | 1.12E+08 | + | 1.26E+01 | 1.08E+00 | 7.87E-01 | 0.008141 | 8.45E-01 | 4.18E-01 | -2.39E+00 | 7.41E-01 | 1.77E+00 | -4.09E+00 | -1.40E+00 |
| Gpc2 | 5 | 1.38E+08 | 1.38E+08 | - | 6.44E+00 | 5.50E-01 | 7.87E-01 | 0.008311 | 8.45E-01 | 4.13E-01 | -2.42E+00 | 3.76E-01 | 9.11E-01 | -4.44E+00 | -1.32E+00 |
| Zfp454 | 11 | 50872723 | 50887502 | - | 1.47E+00 | 1.27E-01 | 7.19E-01 | 0.002750 | 7.73E-01 | 3.96E-01 | -2.52E+00 | 8.37E-02 | 2.11E-01 | -4.22E+00 | -1.51E+00 |
| Rgs9 | 11 | 1.09E+08 | 1.09E+08 | - | 8.47E-01 | 7.21E-02 | 7.87E-01 | 0.008125 | 8.45E-01 | 3.89E-01 | -2.57E+00 | 4.74E-02 | 1.22E-01 | -4.49E+00 | -1.47E+00 |
| Azgp1 | 5 | 1.38E+08 | 1.38E+08 | + | 5.36E-01 | 4.55E-02 | 7.19E-01 | 0.003506 | 7.73E-01 | 3.86E-01 | -2.59E+00 | 2.98E-02 | 7.73E-02 | -4.43E+00 | -1.52E+00 |
| Gm11110 | 17 | 57092023 | 57105943 | - | 7.21E-01 | 6.13E-02 | 7.19E-01 | 0.003652 | 7.73E-01 | 3.81E-01 | -2.62E+00 | 3.98E-02 | 1.04E-01 | -4.45E+00 | -1.55E+00 |
| Tubb4a | 17 | 57080066 | 57087783 | - | 1.58E+00 | 1.34E-01 | 7.19E-01 | 0.002563 | 7.73E-01 | 3.81E-01 | -2.62E+00 | 8.70E-02 | 2.28E-01 | -4.29E+00 | -1.60E+00 |
| Rtl6 | 15 | 84553398 | 84557824 | - | 1.51E+01 | 1.28E+00 | 7.74E-01 | 0.004622 | 8.31E-01 | 3.79E-01 | -2.64E+00 | 8.33E-01 | 2.20E+00 | -4.50E+00 | -1.54E+00 |
| Arg2 | 12 | 79130788 | 79156302 | + | 5.46E+00 | 4.65E-01 | 7.19E-01 | 0.002896 | 7.73E-01 | 3.76E-01 | -2.66E+00 | 2.98E-01 | 7.93E-01 | -4.41E+00 | -1.61E+00 |
| LOC100861615 | 14 | 6259747 | 6287251 | - | 4.52E-01 | 3.73E-02 | 8.23E-01 | 0.009781 | 8.84E-01 | 3.64E-01 | -2.75E+00 | 2.41E-02 | 6.63E-02 | -5.16E+00 | -1.46E+00 |
| Ankrd33b | 15 | 31291479 | 31367760 | - | 5.94E-01 | 4.81E-02 | 7.96E-01 | 0.008952 | 8.56E-01 | 3.44E-01 | -2.90E+00 | 3.04E-02 | 8.83E-02 | -5.65E+00 | -1.49E+00 |
| Morn4 | 19 | 42074939 | 42086371 | - | 3.09E+00 | 2.53E-01 | 7.19E-01 | 0.003181 | 7.73E-01 | 3.36E-01 | -2.98E+00 | 1.55E-01 | 4.62E-01 | -5.40E+00 | -1.64E+00 |
| Fkbp6 | 5 | 1.35E+08 | 1.35E+08 | - | 6.81E-02 | 5.21E-03 | 7.87E-01 | 0.007610 | 8.45E-01 | 3.05E-01 | -3.28E+00 | 3.18E-03 | 1.04E-02 | -6.74E+00 | -1.60E+00 |
| 1700010I14Rik | 17 | 8988333 | 9008320 | + | 7.10E-01 | 5.24E-02 | 7.87E-01 | 0.007181 | 8.45E-01 | 3.04E-01 | -3.29E+00 | 3.31E-02 | 1.09E-01 | -7.51E+00 | -1.44E+00 |
| Panx2 | 15 | 89059726 | 89070908 | + | 7.20E-01 | 5.46E-02 | 7.19E-01 | 0.001701 | 7.73E-01 | 3.02E-01 | -3.31E+00 | 3.34E-02 | 1.11E-01 | -6.03E+00 | -1.82E+00 |
| Myo1e | 9 | 70207350 | 70400068 | + | 3.01E+00 | 2.46E-01 | 1.69E-02 | 0.000003 | 1.82E-02 | 2.81E-01 | -3.57E+00 | 1.32E-01 | 4.70E-01 | -4.60E+00 | -2.76E+00 |
| D630003M21Rik | 2 | 1.58E+08 | 1.58E+08 | - | 2.72E-01 | 1.88E-02 | 7.87E-01 | 0.007559 | 8.45E-01 | 2.74E-01 | -3.64E+00 | 1.17E-02 | 4.26E-02 | -9.43E+00 | -1.41E+00 |
| Nlgn3 | X | 1.01E+08 | 1.01E+08 | + | 2.70E-01 | 1.93E-02 | 7.87E-01 | 0.008163 | 8.45E-01 | 2.65E-01 | -3.77E+00 | 1.13E-02 | 4.27E-02 | -8.55E+00 | -1.66E+00 |
| 6820408C15Rik | 2 | 1.52E+08 | 1.52E+08 | + | 1.13E-01 | 8.06E-03 | 7.87E-01 | 0.006099 | 8.45E-01 | 2.57E-01 | -3.90E+00 | 4.63E-03 | 1.81E-02 | -9.71E+00 | -1.56E+00 |
| Acsbg1 | 9 | 54604997 | 54661886 | - | 9.88E-01 | 6.95E-02 | 7.87E-01 | 0.006959 | 8.45E-01 | 2.51E-01 | -3.98E+00 | 3.97E-02 | 1.58E-01 | -8.96E+00 | -1.77E+00 |
| Catsperd | 17 | 56627047 | 56664457 | + | 3.93E-01 | 2.74E-02 | 7.84E-01 | 0.005532 | 8.43E-01 | 2.43E-01 | -4.12E+00 | 1.53E-02 | 6.32E-02 | -9.15E+00 | -1.85E+00 |
| Drd4 | 7 | 1.41E+08 | 1.41E+08 | + | 9.38E-01 | 6.39E-02 | 7.87E-01 | 0.008402 | 8.45E-01 | 2.36E-01 | -4.24E+00 | 3.58E-02 | 1.52E-01 | -1.18E+01 | -1.52E+00 |
| Mtmr7 | 8 | 40550159 | 40634793 | - | 2.75E+00 | 1.91E-01 | 7.19E-01 | 0.002730 | 7.73E-01 | 2.31E-01 | -4.32E+00 | 1.03E-01 | 4.47E-01 | -9.71E+00 | -1.92E+00 |
| Mns1 | 9 | 72438529 | 72458577 | + | 2.35E+00 | 1.63E-01 | 6.75E-01 | 0.000342 | 7.25E-01 | 1.91E-01 | -5.24E+00 | 7.52E-02 | 3.94E-01 | -9.97E+00 | -2.76E+00 |
| 4930555G01Rik | 14 | 5050629 | 5059244 | + | 3.75E-01 | 2.31E-02 | 7.33E-01 | 0.004066 | 7.87E-01 | 1.85E-01 | -5.39E+00 | 1.17E-02 | 6.33E-02 | -1.34E+01 | -2.17E+00 |
| Wnt4 | 4 | 1.37E+08 | 1.37E+08 | + | 6.21E+00 | 3.99E-01 | 7.19E-01 | 0.001441 | 7.73E-01 | 1.78E-01 | -5.61E+00 | 1.88E-01 | 1.05E+00 | -1.23E+01 | -2.57E+00 |
| Slc35f3 | 8 | 1.26E+08 | 1.26E+08 | + | 6.24E-01 | 3.54E-02 | 7.79E-01 | 0.005193 | 8.37E-01 | 1.58E-01 | -6.31E+00 | 1.71E-02 | 1.08E-01 | -1.95E+01 | -2.04E+00 |
| Igsf9b | 9 | 27299104 | 27357550 | + | 6.14E-01 | 3.46E-02 | 6.75E-01 | 0.000685 | 7.25E-01 | 1.32E-01 | -7.57E+00 | 1.43E-02 | 1.09E-01 | -1.89E+01 | -3.03E+00 |
| Kcnh2 | 5 | 24319589 | 24351605 | - | 3.22E-01 | 1.78E-02 | 7.19E-01 | 0.001106 | 7.73E-01 | 1.29E-01 | -7.73E+00 | 7.37E-03 | 5.70E-02 | -1.95E+01 | -3.06E+00 |
| Tmprss6 | 15 | 78439667 | 78468635 | - | 7.84E-01 | 3.68E-02 | 7.19E-01 | 0.001125 | 7.73E-01 | 1.05E-01 | -9.50E+00 | 1.49E-02 | 1.42E-01 | -2.88E+01 | -3.14E+00 |
| Nrxn2 | 19 | 6418738 | 6533218 | + | 3.33E-02 | 1.68E-03 | 6.75E-01 | 0.000496 | 7.25E-01 | 1.05E-01 | -9.52E+00 | 6.32E-04 | 6.02E-03 | -2.26E+01 | -4.02E+00 |
| Col6a6 | 9 | 1.06E+08 | 1.06E+08 | - | 3.58E-01 | 1.40E-02 | 7.87E-01 | 0.007099 | 8.45E-01 | 9.22E-02 | -1.08E+01 | 6.05E-03 | 6.56E-02 | -4.90E+01 | -2.40E+00 |
| Adra1a | 14 | 66635251 | 66771169 | + | 1.30E-01 | 4.39E-03 | 7.74E-01 | 0.004755 | 8.31E-01 | 7.80E-02 | -1.28E+01 | 1.88E-03 | 2.40E-02 | -6.36E+01 | -2.59E+00 |
| Amtn | 5 | 88376108 | 88385917 | + | 9.73E+00 | 3.00E-01 | 7.87E-01 | 0.006787 | 8.45E-01 | 6.35E-02 | -1.57E+01 | 1.16E-01 | 1.83E+00 | -7.77E+01 | -3.19E+00 |
| Clca1 | 3 | 1.45E+08 | 1.45E+08 | - | 1.51E+00 | 4.91E-02 | 7.87E-01 | 0.006673 | 8.45E-01 | 6.06E-02 | -1.65E+01 | 1.72E-02 | 2.84E-01 | -6.90E+01 | -3.95E+00 |

Supplemental Table 1) **Gene set analysis results**. Gene set analysis results of RNA-seq data using Partek® Flow software. The analysis of RNA-seq data from Myo1e WT and Myo1e KO tumors yielded 79 differentially expressed genes beyond +/- 1.5 fold expression change and a relative expression p-value of $p<0.01$.

| Master Regulator | Molecule Type | Depth | Predicted Activation State | Notes | Activation z-score | p-value of overlap | Network bias-corrected p-value | Target molecules in dataset |
|---|---|---|---|---|---|---|---|---|
| POSTN | other | 3 | | | -1.342 | 1.86E-03 | 5.20E-03 | ADRA1A,ARG2,AZGP1,CLCA1,DUSP14,FKBP6,Gimap9,IGSF9B,KCNH2,LALBA,MNS1,MTMR7,MYH10,MYO1E,NR2F2,OSGIN1, PIGR,PSMC3IP,RASSF1,TMPRSS6 |
| Mlc | group | 3 | | | -0.775 | 2.38E-03 | 4.70E-03 | ACSBG1,ADRA1A,ARG2,AZGP1,DUSP14,IGSF9B,KCNH2,MNS1,MTMR7,MYO1E,NR2F2,PHACTR1,PIGR,RASSF1,TUBB4A |
| ERK | group | 3 | | | -1.461 | 2.43E-03 | 8.80E-03 | ACSBG1,ADRA1A,Apol7e (includes others),ARG2,CLCA1,DRD4,DUSP14,Fcna,FKBP6,Gimap9,IGSF9B,KCNH2,LPCAT4,MNS1, MRGPRE,MYO1E,NLGN3,NR2F2,NRXN2,OSGIN1,PIGR,Podxl,PSMC3IP,RASSF1,STEAP1,TLN2,TMEM2,TUBB4A,VSIG4,XK |
| PANX3 | transporter | 2 | | | 0.378 | 2.73E-03 | 1.60E-03 | ADRA1A,DUSP14,Gimap9,IGSF9B,MNS1,MYH10,PSMC3IP |
| GPBAR1 | G-protein coupled receptor | 3 | | | -1 | 2.75E-03 | 1.60E-03 | ADRA1A,ARG2,DUSP14,FKBP6,Gimap9,IGSF9B,KCNH2,LALBA,MNS1,MTMR7,MYH10,NR2F2,OSGIN1,PIGR,PSMC3IP,WNT4 |
| MAP2 | other | 2 | | | -0.378 | 2.78E-03 | 1.70E-03 | ADRA1A,DUSP14,Gimap9,IGSF9B,MNS1,MYH10,PSMC3IP |
| HLA-B | transmembrane receptor | 3 | | | 0.775 | 3.02E-03 | 1.60E-03 | Apol7e (includes others),AZGP1,DUSP14,KCNH2,LALBA,MNS1,NR2F2,OSGIN1,PHACTR1,PIGR,Podxl,RASSF1,TMPRSS6,TUBB4A, WNT4 |
| RASSF1-AS1 | other | 1 | | biased | -1 | 3.15E-03 | 3.00E-03 | RASSF1 |
| miR-602 (miRNAs w/seed ACACGGG) | mature microRNA | 1 | | biased | -1 | 3.15E-03 | 3.00E-03 | RASSF1 |
| mir-602 | microRNA | 1 | | biased | -1 | 3.15E-03 | 3.00E-03 | RASSF1 |
| EAF1 | transcription regulator | 1 | | biased | 1 | 3.15E-03 | 3.30E-03 | WNT4 |
| mir-584 | microRNA | 1 | | biased | 1 | 3.15E-03 | 3.60E-03 | PHACTR1 |
| RGS9BP | other | 1 | | biased | -1 | 3.15E-03 | 4.30E-03 | RGS9 |
| RNF207 | other | 1 | | biased | -1 | 3.15E-03 | 6.20E-03 | KCNH2 |
| GP1BA | transmembrane receptor | 3 | | | -0.943 | 3.58E-03 | 5.40E-03 | ADRA1A,ARG2,AZGP1,DUSP14,FKBP6,IGSF9B,KCNH2,LALBA,MNS1,MYH10,MYO1E,OSGIN1,PHACTR1,PIGR,PSMC3IP,RASSF1,TUBB4A,WNT4 |
| LGMN | peptidase | 3 | | | -1.508 | 4.12E-03 | 1.20E-03 | ARG2,Gimap9,KCNH2,LALBA,MNS1,MTMR7,MYH10,PIGR,PSMC3IP,TUBB4A,WNT4 |
| Camk | complex | 2 | | | -0.707 | 4.96E-03 | 1.10E-03 | ADRA1A,DUSP14,Gimap9,IGSF9B,KCNH2,MNS1,MYH10,PSMC3IP |
| PTK2 | kinase | 2 | | | -0.728 | 5.31E-03 | 6.40E-03 | ADRA1A,ARG2,AZGP1,DUSP14,FKBP6,IGSF9B,KCNH2,LALBA,MNS1,MYH10,MYO1E,OSGIN1,PHACTR1,PIGR,RASSF1,TUBB4A,WNT4 |
| JAG1 | growth factor | 1 | | biased | 1.414 | 5.45E-03 | 1.90E-03 | NR2F2,WNT4 |
| PDCL | enzyme | 1 | | biased | -1 | 6.29E-03 | 8.50E-03 | RGS9 |
| SIGMAR1 | transmembrane receptor | 1 | | biased | -1 | 6.29E-03 | 9.40E-03 | KCNH2 |
| ADORA2A | G-protein coupled receptor | 2 | | | -0.577 | 6.36E-03 | 1.90E-03 | ADRA1A,ARG2,AZGP1,DUSP14,Gimap9,IGSF9B,KCNH2,MNS1,MYH10,PIGR,PSMC3IP,TUBB4A |
| cytochrome C | group | 3 | | | -1 | 6.37E-03 | 1.22E-02 | ADRA1A,ARG2,AZGP1,CLCA1,DUSP14,FKBP6,Gimap9,IGSF9B,KCNH2,LALBA,MNS1,OSGIN1,PHACTR1,PIGR,PSMC3IP,WNT4 |
| FHL3 | other | 2 | | | -1.134 | 6.61E-03 | 2.90E-03 | ADRA1A,DUSP14,IGSF9B,MNS1,MYH10,RASSF1,TMPRSS6 |
| RPS6KA5 | kinase | 2 | | biased | -0.577 | 6.99E-03 | 1.50E-03 | ADRA1A,ARG2,AZGP1,DUSP14,FKBP6,Gimap9,IGSF9B,MNS1,MYH10,PHACTR1,PSMC3IP,XK |
| EPAS1 | transcription regulator | 1 | | biased | -0.577 | 7.08E-03 | 1.60E-03 | ARG2,RASSF1,TMPRSS6 |
| GNAS | enzyme | 2 | | | -0.632 | 7.99E-03 | 2.40E-03 | ADRA1A,ARG2,DUSP14,Gimap9,IGSF9B,KCNH2,MNS1,MYH10,PIGR,PSMC3IP |
| RASA2 | other | 2 | | biased | 1.633 | 8.16E-03 | 2.90E-03 | ADRA1A,DUSP14,IGSF9B,MNS1,MYH10,PIGR |
| PASK | kinase | 2 | | | -0.378 | 8.35E-03 | 2.10E-03 | ADRA1A,DUSP14,IGSF9B,KCNH2,MNS1,MYH10,TUBB4A |
| MAP2K2 | kinase | 2 | | | -1.414 | 9.17E-03 | 9.00E-04 | ACSBG1,ARG2,Gimap9,KCNH2,MNS1,MYH10,PIGR,PSMC3IP |
| EAF2 | transcription regulator | 1 | | | 1 | 9.43E-03 | 7.50E-03 | WNT4 |
| WNT9B | other | 1 | | biased | -1 | 9.43E-03 | 9.40E-03 | WNT4 |
| GPR17 | G-protein coupled receptor | 3 | | | 0.229 | 9.45E-03 | 2.69E-02 | ADRA1A,ARG2,AZGP1,CLCA1,DRD4,DUSP14,Gimap9,IGSF9B,KCNH2,LALBA,MNS1,MTMR7,PHACTR1,PIGR,Podxl,PSMC3IP,STEAP1,TUBB4A, VSIG4 |
| BCL2A1 | other | 3 | | | -1.155 | 9.82E-03 | 1.06E-02 | AZGP1,CLCA1,DUSP14,Gimap9,KCNH2,LALBA,MNS1,MYH10,NR2F2,PIGR,PSMC3IP,WNT4 |
| LYPD6 | other | 3 | | | -0.243 | 9.94E-03 | 2.28E-02 | ACSBG1,ADRA1A,DUSP14,IGSF9B,LALBA,LPCAT4,MNS1,MTMR7,PHACTR1,PIGR,PSMC3IP,RASSF1,STEAP1,TMEM2,TMPRSS6,TUBB4A,WNT4 |
| WNT2 | cytokine | 3 | | | -0.243 | 1.00E-02 | 2.29E-02 | ACSBG1,ADRA1A,DUSP14,IGSF9B,LALBA,LPCAT4,MNS1,MTMR7,PHACTR1,PIGR,PSMC3IP,RASSF1,STEAP1,TMEM2,TMPRSS6,TUBB4A,WNT4 |
| FZD1 | G-protein coupled receptor | 3 | | | -0.728 | 1.04E-02 | 2.63E-02 | ACSBG1,ADRA1A,DUSP14,IGSF9B,LALBA,LPCAT4,MTMR7,NR2F2,PHACTR1,PIGR,PSMC3IP,RASSF1,STEAP1,TMEM2,TMPRSS6,TUBB4A,WNT4 |
| PKC Îµ/Î´ | group | 3 | | | 1.342 | 1.10E-02 | 3.23E-02 | ADRA1A,AZGP1,DUSP14,FKBP6,Gimap9,IGSF9B,KCNH2,LPCAT4,MNS1,MTMR7,NR2F2,NRXN2,OSGIN1,PHACTR1,PSMC3IP,PXYLP1,STEAP1, TUBB4A, WNT4,XK |
| RPS6KA2 | kinase | 3 | | | -0.943 | 1.13E-02 | 1.96E-02 | ACSBG1,ADRA1A,AZGP1,DUSP14,FKBP6,IGSF9B,KCNH2,LALBA,MNS1,MTMR7,MYH10,NR2F2,PHACTR1,PIGR,RASSF1,STEAP1,TMPRSS6,TUBB4A |
| SIK2 | kinase | 3 | Inhibited | | -2 | 1.17E-02 | 2.14E-02 | ACSBG1,ADRA1A,AZGP1,DUSP14,Gimap9,IGSF9B,LALBA,MTMR7,MYH10,NR2F2,PHACTR1,PIGR,PSMC3IP,RASSF1,TMPRSS6,TUBB4A |
| LRP | group | 3 | Inhibited | | -2.111 | 1.19E-02 | 9.00E-03 | ACSBG1,ADRA1A,DUSP14,IGSF9B,KCNH2,MNS1,MYH10,NR2F2,PHACTR1,TMEM2,TUBB4A |
| ULK1 | kinase | 1 | | | -1 | 1.25E-02 | 9.70E-03 | TUBB4A |
| RAB4A | enzyme | 1 | | biased | 1 | 1.25E-02 | 1.55E-02 | KCNH2 |
| RS1 | other | 1 | | biased | 1 | 1.25E-02 | 1.73E-02 | RGS9 |
| Erm | group | 3 | | | 1.291 | 1.30E-02 | 3.07E-02 | ACSBG1,ADRA1A,AZGP1,CLCA1,DRD4,DUSP14,Gimap9,IGSF9B,PHACTR1,Podxl,PSMC3IP,RASSF1,TMPRSS6,TUBB4A,VSIG4 |
| Calmodulin | group | 2 | | | -0.577 | 1.30E-02 | 5.80E-03 | ADRA1A,ARG2,AZGP1,DRD4,DUSP14,Gimap9,IGSF9B,KCNH2,MNS1,MYH10,PIGR,PSMC3IP |
| CREB1 | transcription regulator | 1 | | | -1.342 | 1.30E-02 | 9.20E-03 | ADRA1A,DUSP14,IGSF9B,MNS1,MYH10 |
| UBQLN1 | other | 3 | | | 0.688 | 1.35E-02 | 2.59E-02 | Apol7e (includes others),ARG2,AZGP1,CLCA1,DUSP14,FKBP6,Gimap9,KCNH2,LALBA,MNS1,MYH10,NR2F2,OSGIN1,PHACTR1,PIGR,PSMC3IP, RASSF1, TMPRSS6,TUBB4A |
| MAPT | other | 2 | | | -1.134 | 1.45E-02 | 2.00E-03 | ARG2,Gimap9,MTMR7,PIGR,PSMC3IP,TUBB4A,WNT4 |
| SV2C | other | 3 | | | -0.655 | 1.53E-02 | 4.95E-02 | ACSBG1,ADRA1A,Apol7e (includes others),ARG2,AZGP1,CLCA1,FKBP6,Gimap9,KCNH2,LALBA,MNS1,MTMR7,NRXN2,OSGIN1,PHACTR1, PSMC3IP, RASSF1,STEAP1, TMPRSS6, TUBB4A,XK |
| PROKR1 | G-protein coupled receptor | 1 | | biased | 1 | 1.57E-02 | 5.60E-03 | STEAP1 |
| FAM20C | enzyme | 1 | | biased | -1 | 1.57E-02 | 1.44E-02 | AMTN |
| TSC2 | other | 2 | | | 0.333 | 1.57E-02 | 6.20E-03 | ARG2,DRD4,KCNH2,MYH10,PHACTR1,PIGR,RASSF1,TMPRSS6,TUBB4A |
| HOXA10 | transcription regulator | 1 | | | 1.732 | 1.68E-02 | 1.31E-02 | PIGR,RNF227,WNT4 |
| BMP6 | growth factor | 1 | | biased | -1.414 | 1.84E-02 | 4.20E-03 | MYH10,TMPRSS6 |
| Scf | complex | 3 | | | -0.5 | 1.85E-02 | 2.84E-02 | AZGP1,CLCA1,DUSP14,KCNH2,LALBA,MTMR7,NR2F2,PHACTR1,Podxl,PSMC3IP,RGS9,STEAP1,TMPRSS6,TUBB4A,WNT4,XK |
| CSE1L | transporter | 1 | | | -1 | 1.88E-02 | 1.52E-02 | RASSF1 |
| GNB5 | enzyme | 1 | | biased | -1 | 1.88E-02 | 1.61E-02 | RGS9 |
| H1FX | other | 1 | | biased | 1 | 1.88E-02 | 2.07E-02 | AMTN |
| NR2E3 | ligand-dependent nuclear receptor | 1 | | biased | 1 | 1.88E-02 | 2.34E-02 | CLCA1 |
| SOX4 | transcription regulator | 1 | | biased | -1.732 | 1.98E-02 | 2.40E-02 | NRXN2,PXYLP1,TMEM2 |
| CREM | transcription regulator | 2 | | | -1.342 | 2.13E-02 | 3.80E-03 | ARG2,DUSP14,Gimap9,MNS1,PSMC3IP |
| DUSP6 | phosphatase | 1 | | | 1 | 2.19E-02 | 1.09E-02 | ARG2 |
| PCBP1 | translation regulator | 1 | | | 1 | 2.19E-02 | 1.50E-02 | MYH10 |
| SPARC | other | 2 | | | -0.632 | 2.35E-02 | 1.25E-02 | ARG2,KCNH2,MNS1,MTMR7,MYH10,NR2F2,PSMC3IP,RGS9,TUBB4A,WNT4 |
| CDK5R1 | kinase | 2 | | | -1 | 2.41E-02 | 7.30E-03 | ACSBG1,ARG2,Gimap9,MNS1,MTMR7,NR2F2,PSMC3IP,TUBB4A,WNT4 |
| SHANK3 | other | 1 | | biased | -1 | 2.49E-02 | 2.54E-02 | NLGN3 |
| HTR1A | G-protein coupled receptor | 2 | | | 0.816 | 2.73E-02 | 3.60E-03 | ARG2,Gimap9,KCNH2,PIGR,PSMC3IP,TUBB4A |
| TLR3 | transmembrane receptor | 1 | | biased | 0.577 | 2.83E-02 | 5.20E-03 | Apol7e (includes others),ARG2,CLCA1 |
| HSPA1A/HSPA1B | enzyme | 1 | | biased | -1 | 3.11E-02 | 1.09E-02 | KCNH2 |
| CREM | transcription regulator | 1 | | | 0 | 3.55E-02 | 1.54E-02 | DUSP14,MNS1 |
| JAK3 | kinase | 2 | | | 0.333 | 3.64E-02 | 1.69E-02 | ARG2,CLCA1,Gimap9,KCNH2,MYH10,PHACTR1,PSMC3IP,RASSF1,TUBB4A |
| HMGN1 | transcription regulator | 1 | | | -1 | 3.72E-02 | 2.54E-02 | AZGP1 |
| Pkar2 | group | 3 | | | 0.378 | 3.97E-02 | 1.92E-02 | ARG2,Gimap9,KCNH2,MNS1,PIGR,PSMC3IP,TUBB4A |
| LGR4 | transmembrane receptor | 1 | | | -1 | 4.02E-02 | 6.90E-03 | WNT4 |
| ADRA1B | G-protein coupled receptor | 1 | | | -1 | 4.02E-02 | 1.78E-02 | ADRA1A |
| MAP2K2 | kinase | 1 | | biased | -1 | 4.02E-02 | 2.09E-02 | ACSBG1 |
| SNX5 | transporter | 3 | | | -0.707 | 4.12E-02 | 1.19E-02 | ARG2,DRD4,Gimap9,KCNH2,MTMR7,PIGR,PSMC3IP,WNT4 |
| EOMES | transcription regulator | 1 | | | 1.414 | 4.27E-02 | 3.84E-02 | DUSP14,TMEM2 |
| HHEX | transcription regulator | 1 | | | -1 | 4.32E-02 | 2.37E-02 | MYH10 |
| ZFPM2 | transcription regulator | 1 | | biased | 1 | 4.32E-02 | 2.46E-02 | WNT4 |
| COL6A1 | other | 1 | | | -1 | 4.32E-02 | 3.88E-02 | COL6A6 |
| Shc | group | 2 | | biased | -1.667 | 4.46E-02 | 1.28E-02 | ACSBG1,ARG2,KCNH2,MNS1,MYH10,PIGR,RASSF1,TMPRSS6,TUBB4A |
| MS4A1 | other | 3 | | | -0.302 | 4.65E-02 | 4.93E-02 | ARG2,CLCA1,DUSP14,KCNH2,LALBA,MNS1,MYH10,PHACTR1,PIGR,STEAP1,WNT4 |
| DNAJB6 | transcription regulator | 1 | | | 1 | 4.93E-02 | 2.12E-02 | AZGP1 |
| PRMT1 | enzyme | 1 | | biased | -1 | 4.93E-02 | 2.68E-02 | ARG2 |
| IL11RA | transmembrane receptor | 1 | | biased | -1 | 5.23E-02 | 1.29E-02 | CLCA1 |
| UBE3A | enzyme | 1 | | biased | 1 | 5.23E-02 | 2.35E-02 | WNT4 |
| SMYD3 | enzyme | 1 | | biased | -1 | 5.23E-02 | 3.71E-02 | RASSF1 |
| Pld | group | 2 | | biased | -0.816 | 5.38E-02 | 1.35E-02 | ARG2,KCNH2,MNS1,MTMR7,PIGR,WNT4 |
| GCH1 | enzyme | 3 | | biased | 0 | 5.50E-02 | 3.16E-02 | ARG2,AZGP1,CLCA1,KCNH2,OSGIN1,PHACTR1,PIGR,RASSF1,TMPRSS6,WNT4 |
| Hmgn3 | other | 1 | | biased | -1 | 5.53E-02 | 3.76E-02 | AZGP1 |
| PAX2 | transcription regulator | 1 | | biased | -1 | 5.53E-02 | 4.18E-02 | WNT4 |
| GHRHR | G-protein coupled receptor | 2 | | biased | 0 | 5.76E-02 | 7.10E-03 | Gimap9,KCNH2,PIGR,PSMC3IP |
| RUNX1 | transcription regulator | 1 | | | 1.414 | 5.92E-02 | 2.62E-02 | MYH10,NR2F2 |
| TREM1 | transmembrane receptor | 2 | | biased | -0.816 | 6.09E-02 | 1.05E-02 | ARG2,DUSP14,KCNH2,MNS1,OSGIN1,PIGR |
| RBM3 | other | 3 | | biased | -0.632 | 6.10E-02 | 3.27E-02 | Apol7e (includes others),ARG2,DUSP14,LALBA,NR2F2,OSGIN1,PHACTR1,RASSF1,TUBB4A,WNT4 |
| POSTN | other | 1 | | biased | 1 | 6.12E-02 | 2.15E-02 | CLCA1 |
| Dynamin | group | 2 | | biased | -1.134 | 6.29E-02 | 2.22E-02 | ARG2,AZGP1,Gimap9,KCNH2,MYH10,PIGR,PSMC3IP |
| Relaxin | group | 1 | | | 1 | 6.71E-02 | 2.02E-02 | ARG2 |
| HEY1 | transcription regulator | 1 | | biased | 1 | 6.71E-02 | 3.22E-02 | NR2F2 |
| ANGPT2 | growth factor | 2 | | | -0.302 | 6.72E-02 | 4.90E-02 | ARG2,AZGP1,DUSP14,Gimap9,KCNH2,MNS1,MTMR7,NR2F2,PIGR,PSMC3IP,WNT4 |
| COL4A4 | other | 3 | | biased | 0 | 6.93E-02 | 3.63E-02 | Apol7e (includes others),ARG2,DUSP14,Gimap9,OSGIN1,PHACTR1,PSMC3IP,RASSF1,TUBB4A,WNT4 |
| TNFSF11 | cytokine | 2 | | biased | -1.508 | 6.96E-02 | 4.85E-02 | ADRA1A,ARG2,AZGP1,KCNH2,MNS1,MYH10,PHACTR1,PIGR,PSMC3IP,TUBB4A,WNT4 |
| CLEC7A | transmembrane receptor | 1 | | biased | -1 | 7.01E-02 | 6.90E-03 | CLCA1 |
| SPRY2 | other | 2 | | | 0.707 | 7.01E-02 | 4.02E-02 | ADRA1A,ARG2,KCNH2,LALBA,MYH10,PIGR,RASSF1,TMPRSS6 |
| GPR183 | G-protein coupled receptor | 2 | | | 1 | 7.05E-02 | 1.77E-02 | ARG2,Gimap9,KCNH2,PSMC3IP |
| BDKRB1 | G-protein coupled receptor | 2 | | biased | -1 | 7.72E-02 | 9.20E-03 | ARG2,AZGP1,KCNH2,PIGR |
| DLG4 | kinase | 2 | | | 0 | 7.76E-02 | 1.65E-02 | ARG2,KCNH2,LALBA,TUBB4A |
| COMT | enzyme | 2 | | | 0.577 | 7.79E-02 | 3.53E-02 | DRD4,PIGR,RASSF1 |
| CSL-HIF1A-MAML1-NICD | complex | 2 | | | 0 | 8.02E-02 | 1.90E-02 | NR2F2,RASSF1,TMPRSS6,WNT4 |
| NRP2 | kinase | 2 | | biased | -0.447 | 8.23E-02 | 3.34E-02 | ARG2,MNS1,MTMR7,MYH10,WNT4 |
| IL6ST | transmembrane receptor | 2 | | | -1.414 | 8.23E-02 | 4.97E-02 | ARG2,KCNH2,MNS1,NR2F2,PHACTR1,PSMC3IP,RASSF1,WNT4 |
| CLDN4 | transmembrane receptor | 3 | | | -0.378 | 8.27E-02 | 3.15E-02 | ARG2,KCNH2,LALBA,MNS1,MYH10,PIGR,WNT4 |
| FHL1 | other | 2 | | | 0 | 8.29E-02 | 2.04E-02 | NR2F2,RASSF1,TMPRSS6,WNT4 |
| CLDN1 | other | 3 | | | -0.378 | 8.30E-02 | 3.16E-02 | ARG2,KCNH2,LALBA,MNS1,MYH10,PIGR,WNT4 |
| NRL | transcription regulator | 3 | | | 0.378 | 8.30E-02 | 4.24E-02 | CLCA1,Fcna,Gimap9,KCNH2,NRXN2,PSMC3IP,VSIG4 |
| SCD | enzyme | 2 | | biased | 0 | 8.44E-02 | 2.29E-02 | ADRA1A,ARG2,AZGP1,MYH10,PHACTR1,PIGR |
| WT1 | transcription regulator | 1 | | | 0 | 8.50E-02 | 4.27E-02 | Podxl,WNT4 |
| GST | complex | 3 | | | -0.378 | 8.56E-02 | 3.78E-02 | ARG2,KCNH2,MNS1,NR2F2,PHACTR1,PIGR,WNT4 |
| Hdac | group | 1 | | | 0 | 8.97E-02 | 4.39E-02 | KCNH2,WNT4 |
| PPP1R17 | other | 2 | | | -1 | 9.02E-02 | 3.54E-02 | ARG2,MNS1,MYH10,TUBB4A |
| YAP1 | transcription regulator | 2 | | biased | -0.816 | 9.30E-02 | 2.75E-02 | ARG2,KCNH2,LALBA,MYH10,NR2F2,WNT4 |
| KIT | transmembrane receptor | 2 | | biased | -1.134 | 9.36E-02 | 2.75E-02 | ARG2,KCNH2,MNS1,MYH10,PHACTR1,PIGR,WNT4 |
| MC4R | G-protein coupled receptor | 2 | | | -1 | 9.64E-02 | 1.66E-02 | ARG2,KCNH2,MYH10,PIGR |
| Hnf3 | group | 3 | | | 1 | 9.78E-02 | 3.97E-02 | ARG2,CLCA1,NR2F2,RASSF1 |
| NOS2 | enzyme | 1 | | | 0 | 9.85E-02 | 3.29E-02 | AZGP1,PIGR |
| IRAK3 | kinase | 2 | | | 1.633 | 1.02E-01 | 2.97E-02 | ARG2,AZGP1,KCNH2,MNS1,PIGR,WNT4 |
| UCN (family) | group | 3 | | biased | -1.134 | 1.03E-01 | 3.68E-02 | ARG2,Gimap9,KCNH2,MNS1,NR2F2,PIGR,PSMC3IP |
| GSKIP | other | 2 | | biased | 1.732 | 1.03E-01 | 4.14E-02 | MTMR7,TUBB4A,WNT4 |
| GIPC1 | other | 2 | Activated | | 2 | 1.04E-01 | 2.50E-02 | ARG2,KCNH2,MNS1,PIGR |
| GSTP1 | enzyme | 2 | | | 0 | 1.06E-01 | 4.58E-02 | ARG2,KCNH2,MNS1,PHACTR1,PIGR,WNT4 |
| PP1 protein complex group | complex | 2 | | | 0 | 1.07E-01 | 3.45E-02 | ARG2,Gimap9,MNS1,PIGR,PSMC3IP,TUBB4A |
| DDX3X | enzyme | 3 | | | 0.447 | 1.09E-01 | 3.32E-02 | ARG2,KCNH2,MTMR7,TUBB4A,WNT4 |
| UCN3 | other | 2 | | biased | 0 | 1.11E-01 | 2.19E-02 | ARG2,Gimap9,KCNH2,PSMC3IP |
| MAML1 | transcription regulator | 2 | | | -0.447 | 1.11E-01 | 3.35E-02 | AZGP1,KCNH2,MYH10,NR2F2,WNT4 |
| Hif1 | complex | 2 | | biased | -0.447 | 1.11E-01 | 4.56E-02 | NR2F2,PSMC3IP,RASSF1,RGS9,TMPRSS6 |
| ADRA2B | G-protein coupled receptor | 2 | Inhibited | | -2 | 1.12E-01 | 2.74E-02 | ARG2,KCNH2,MNS1,PIGR |
| LYPD6 | other | 2 | | biased | -0.577 | 1.15E-01 | 4.40E-02 | MTMR7,TUBB4A,WNT4 |
| WNT2 | cytokine | 2 | | biased | -0.577 | 1.16E-01 | 4.44E-02 | MTMR7,TUBB4A,WNT4 |
| NEK10 | kinase | 2 | | biased | -1.414 | 1.18E-01 | 1.69E-02 | ACSBG1,PIGR |
| GADD45G | other | 2 | | biased | -0.447 | 1.18E-01 | 3.82E-02 | ARG2,AZGP1,PIGR,RASSF1,WNT4 |
| HBEGF | growth factor | 1 | | biased | 1 | 1.19E-01 | 2.83E-02 | LALBA |
| KLRC2 | transmembrane receptor | 2 | | | -1 | 1.19E-01 | 3.18E-02 | ARG2,KCNH2,MNS1,WNT4 |
| PDE10A | enzyme | 2 | | biased | 0.577 | 1.21E-01 | 3.66E-02 | ARG2,Gimap9,PSMC3IP |
| PRKACA | kinase | 1 | | biased | -1 | 1.22E-01 | 4.13E-02 | MYH10 |
| NUMBL | other | 2 | | | 1 | 1.22E-01 | 3.73E-02 | AZGP1,KCNH2,NR2F2,WNT4 |
| CCL19 | cytokine | 2 | | | -0.447 | 1.25E-01 | 3.40E-02 | KCNH2,MYH10,PIGR,TUBB4A,WNT4 |
| RGS3 | other | 2 | Activated | biased | 2.449 | 1.26E-01 | 2.78E-02 | ARG2,KCNH2,MTMR7,MYH10,PIGR,WNT4 |
| PTN | growth factor | 2 | | biased | -0.816 | 1.27E-01 | 4.22E-02 | KCNH2,LALBA,MTMR7,PIGR,TUBB4A,WNT4 |
| Scf Trcp beta | complex | 2 | Activated | biased | 2 | 1.27E-01 | 4.40E-02 | AZGP1,KCNH2,MTMR7,WNT4 |
| IFNB1 | cytokine | 1 | | biased | 1.414 | 1.33E-01 | 4.94E-02 | Apol7e (includes others),RASSF1 |
| STRAP | other | 2 | | | -1.414 | 1.34E-01 | 3.31E-02 | PIGR,WNT4 |
| GLRX | enzyme | 2 | | biased | -0.577 | 1.38E-01 | 4.64E-02 | ADRA1A,CLCA1,WNT4 |
| USP15 | peptidase | 2 | Inhibited | biased | -2 | 1.40E-01 | 4.75E-02 | AZGP1,KCNH2,MTMR7,WNT4 |
| LRP6 | transmembrane receptor | 2 | | | -1 | 1.44E-01 | 4.42E-02 | MTMR7,PHACTR1,TUBB4A,WNT4 |
| NOTCH1 | transcription regulator | 1 | | | 0 | 1.46E-01 | 4.20E-02 | NR2F2,WNT4 |
| CAT | enzyme | 1 | | biased | -1 | 1.49E-01 | 1.71E-02 | CLCA1 |
| MYZAP | other | 2 | | | 0 | 1.50E-01 | 4.91E-02 | ARG2,MTMR7,TUBB4A,WNT4 |
| THBD | transmembrane receptor | 2 | | biased | 0.577 | 1.52E-01 | 4.46E-02 | ARG2,KCNH2,PIGR |
| PAQR3 | other | 2 | | biased | 1 | 1.53E-01 | 2.90E-02 | ARG2,PIGR,RASSF1,TMPRSS6 |
| PLCE1 | enzyme | 2 | | | -1.342 | 1.56E-01 | 4.65E-02 | ARG2,KCNH2,MNS1,MYH10,PIGR |
| DUSP16 | phosphatase | 2 | | | 1 | 1.64E-01 | 4.84E-02 | ARG2,MNS1,PIGR,WNT4 |
| Rasgrf | group | 2 | | biased | 0.577 | 1.73E-01 | 4.91E-02 | MYH10,PIGR,WNT4 |
| LRP5 | transmembrane receptor | 2 | | biased | -1 | 1.73E-01 | 4.36E-02 | KCNH2,MTMR7,TUBB4A,WNT4 |
| TDGF1 | growth factor | 2 | | | 0 | 1.79E-01 | 4.60E-02 | KCNH2,LALBA,PIGR,TUBB4A |
| RAB1A | enzyme | 2 | | biased | -1.732 | 1.82E-01 | 3.59E-02 | ACSBG1,ARG2,KCNH2 |
| myosin-light-chain kinase | group | 2 | | biased | 1.732 | 1.87E-01 | 4.39E-02 | ACSBG1,ARG2,KCNH2 |
| DOK2 | other | 2 | | biased | 1.732 | 2.09E-01 | 4.88E-02 | ARG2,KCNH2,PIGR |
| TRAT1 | kinase | 2 | | biased | 0.577 | 2.21E-01 | 4.93E-02 | ARG2,KCNH2,WNT4 |

Supplemental Table 2) **Upstream causal network analysis of RNA-seq data**. Ingenuity pathway analysis (IPA, Qiagen Inc., https://www.qiagenbioinformatics.com/products/ingenuity-pathway-analysis) results indicating likely upstream master regulators of the RNA-seq generated gene expression profiles between Myo1e WT and Myo1e KO tumors.
